# A multi-tissue developmental gene expression atlas towards understanding the biological basis of phenotypes in sheep

**DOI:** 10.1101/2024.11.14.623505

**Authors:** Bingru Zhao, Hanpeng Luo, Xuefeng Fu, Guoming Zhang, Emily L. Clark, Feng Wang, Brian Paul Dalrymple, V. Hutton Oddy, Philip E. Vercoe, Cuiling Wu, George E. Liu, Cong-jun Li, Ruidong Xiang, Kechuan Tian, Yanli Zhang, Lingzhao Fang

## Abstract

Sheep (*Ovis aries*) represents one of the most important livestock species for animal protein and wool production worldwide. However, little is known about the genetic and biological basis of ovine phenotypes, particularly for those of high economic value and environmental impact. Here, by generating and integrating 1,413 RNA-seq samples from 51 distinct tissues across 14 developmental time points, representing early prenatal, late prenatal, neonate, lamb, juvenile, adult, and elderly stages, we built a high-resolution developmental Gene Expression Atlas (dGEA) in sheep. We observed dynamic patterns of gene expression and regulatory networks across tissues and developmental stages. When harnessing this resource for interpreting genomic associations of 48 monogenetic and 12 complex traits in sheep, we found that genes upregulated at prenatal developmental stages played more important roles in shaping these phenotypes than those upregulated at postnatal stages. For instance, genetic associations of crimp number, mean staple length (MSL), and individual birth weight were significantly enriched in the prenatal rather than postnatal skin and immune tissues. By comprehensively integrating fine-mapping results and the sheep dGEA, we identified several key genes associated with complex traits in sheep, such as SOX9 (associated with MSL), GNRHR (associated with litter size at birth), and PRKDC (associated with live weight). These results provide novel insights into the gene regulatory and developmental architecture underlying ovine phenotypes. The dGEA (https://sheepdgea.njau.edu.cn/) will serve as an invaluable resource for sheep developmental biology, genetics, genomics, and selective breeding.

## Introduction

Sheep (*Ovis aries*) represent one of the most important domestic animals for meat, milk, and wool production worldwide[1]. To meet the increasing demand for safe animal production, while minimizing the associated negative environmental impacts, a sustainable genetic improvement program is urgently needed in sheep, balancing production, reproduction, health and feed efficiency[2]. A better understanding of the genetic and biological basis of these complex traits will accelerate their current genetic improvement *via* genomic selection and catalyze advanced precision breeding (e.g., genome editing-based) and cell-based food systems in sheep[3–6].

Genome-wide association studies (GWAS) have uncovered many loci contributing to a wide range of complex traits in sheep[7]. For instance, the Animal QTLdb (Aug. 25, 2024) curated 4,743 quantitative trait loci (QTL) from 270 publications, representing 167 base traits and 217 trait variants in sheep[8]. However, the causal genes/variants and molecular mechanisms (e.g., tissues or cell types in which those genomic variants act) underlying most of those complex traits are largely unknown, partially due to 1) high linkage disequilibrium (LD) between causal variants and nearby markers, and 2) the lack of detailed functional annotations of these QTL regions.

Characterization of multi-tissue gene expression could offer valuable insights into the genetic and biological basis of complex traits[9, 10]. In the past decades, great efforts have been made by the scientific community to functionally annotate genes in many species, such as the ENCODE[11] and Genotype-Tissue Expression (GTEx) projects in humans [12–16], and similar initiatives existed for domestic animals including the Functional Annotation of Animal Genomes (FAANG)[16–19] and Farm animal GTEx (FarmGTEx) projects[20, 21]. Since the reference genome was published in 2014[1], several gene expression studies have also been conducted to improve the functional annotation of genes in sheep [5, 22–24]. However, most of these studies are limited to either tissue types or developmental stages[22, 25–28]. A systematic characterization of multiple-tissue developmental transcriptomes will give us a great opportunity to unravel tissues and developmental stages in which trait-associated variants act, providing novel molecular mechanisms underpinning complex traits[29–31].

Here, we built a developmental Gene Expression Atlas (dGEA) in sheep by integrating 410 newly generated RNA-seq samples (30 tissues across 10 developmental time points from 44 animals) and 1,003 publicly available high-quality RNA-seq samples. The dGEA represents a total of 51 distinct tissue/cell types (hereafter referred to as “tissues”) originating from all three germ layers (endoderm, mesoderm, and ectoderm) across 14 developmental time points from day 15 of the embryonic stage (E15) to 7-year-old (Y7, Figure 1A). Based on the known developmental biology in sheep[22], we classified these 14 time points into 7 stages, including early prenatal (≤ E70), late prenatal (>E70), neonatal (D0-D8), lamb (D8-M6), juvenile (M6-Y1.5) adult (Y1.5-Y7), and elderly (>7Y)[22, 32]. We systematically investigated tissue- and development-specific features of the sheep transcriptome, and then utilized these biological insights to explore the developmental and tissue basis of 48 monogenic traits from the Online Mendelian Inheritance in Animals (OMIA) and 12 complex traits with GWAS data (n =1,639) in sheep, including wool production, growth, and reproduction. Our results, for the first time, systemically establish connections among development stage, tissue, and phenotype in sheep. The dGEA (https://sheepdgea.njau.edu.cn/) will serve as a valuable resource for developmental biology, genetics, genomics, and selective breeding in sheep and even other ruminants.

**Figure 1.**
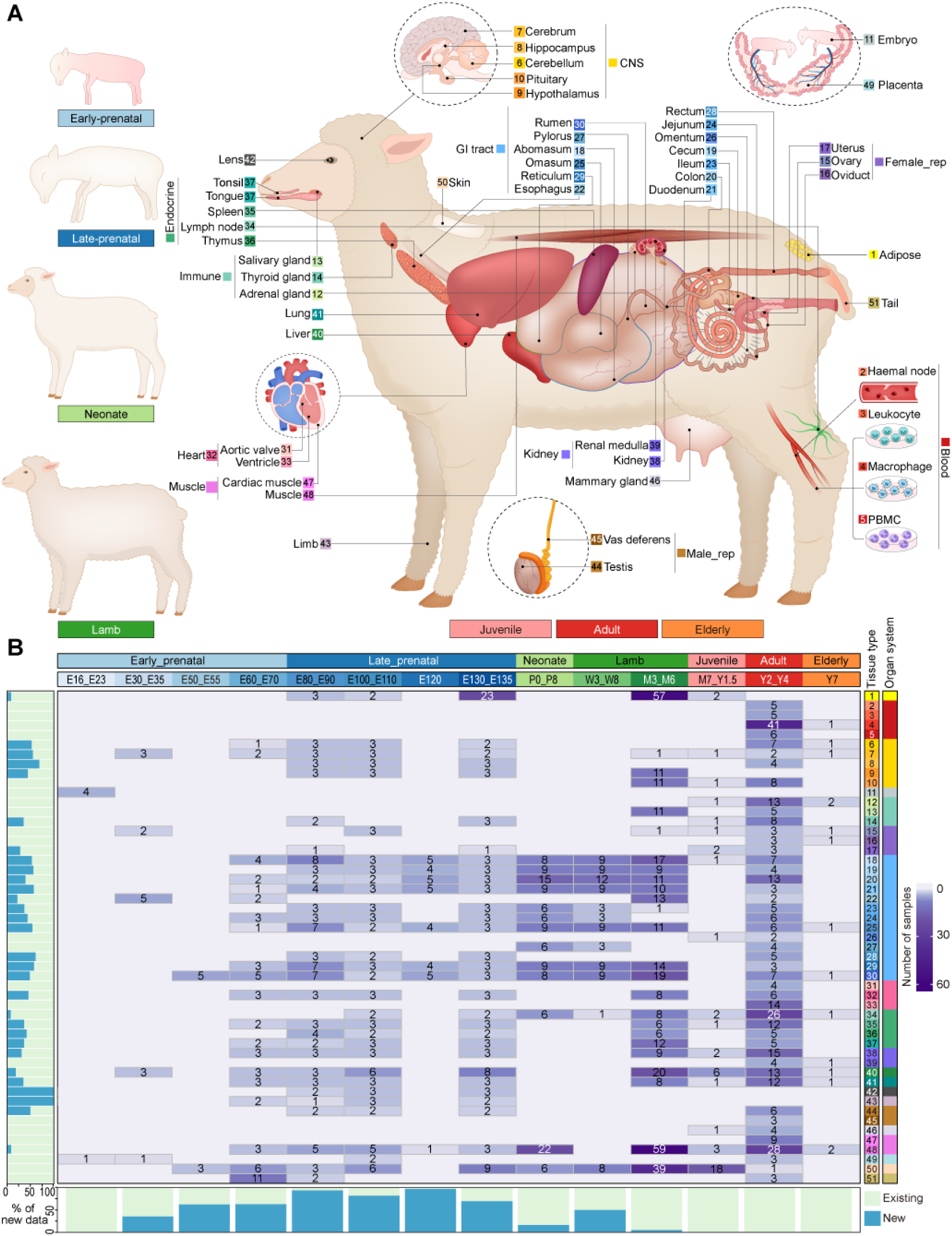
Data summary of the sheep multi-tissue developmental Gene Expression Atlas (dGEA) **A.** Illustration of 51 different tissues and developmental stages in the sheep dGEA, where the color coding for tissues and developmental stages is consistently used in all the figures throughout the entire manuscript. **B.** Distribution of samples from each tissue across various developmental stages. We classified 51 tissues into 20 organ systems (meta-tissue) and 14 time points into 7 developmental stages, i.e., early prenatal (≤ E70), late prenatal (>E70), neonatal (D0-D8), lamb (D8-M6), juvenile (M6-Y1.5) adult (Y1.5-Y7), and elderly (>7Y). The value in the heatmap represents the sample size of a tissue at a developmental stage. The bar charts on the left and bottom represent the proportions of newly generated RNA-seq data in this study across tissues and developmental stages, respectively.

## Results

### Data summary in the sheep developmental Gene Expression Atlas (dGEA)

A total of ~45.5 billion clean reads were yielded with an averaged uniquely mapped rate of 81.79% across all the 1,557 RNA-seq samples (Figure S1; Table S1). After filtering out low-quality samples and non-expressed genes (see Methods), a total of 1,413 RNA-seq samples and 26,423 genes were retained for the subsequent analysis (Figure 1B, Figure S1). Among all the Ensembl annotated genes in the *Ovis aries* reference genome (Oar_rambouillet_v1.0), 26,423 (99.8%) were expressed (Transcripts per million, TPM > 0.1) in at least one sample. The number of expressed genes increased with the number of clean reads across samples and tissues (Figure S2A and B). Among different types of genes (e.g., lncRNA and miRNA), protein-coding genes (PCGs) showed the lowest tissue specificity (TAU value) (Figure S2C), and many of them were also ubiquitously expressed across all the developmental stages (Figure S2D-G).

Hierarchical clustering and principal component analyses (PCA) based on both gene expression and alternative splicing showed a clear separation of RNA-seq samples by tissue type (Figure 2A and B, Figure S3). Within individual tissue types, samples could be clustered according to developmental stages (Figure S4). The most evident separation of samples was between pre- and postnatal stages. When focusing on eight tissues of large sample sizes (n > 15), representing all the developmental stages and three germ layers, we found that in general, expression profiles of prenatal tissues were more like each other compared to postnatal tissues (Figure 2C and D). For instance, prenatal skin samples clustered together with many tissues, while postnatal skin was clearly separated from them (Figure 2C and D). The tissues descended from the same germ layer exhibited more similar gene expression profiles compared to those from different germ layers, confirming that the embryonic origin has a significant influence on tissue-specific gene expression patterns (Figure 2 E-F, Figure S5). All these results support the reliability of this sheep dGEA resource for the subsequent exploration of tissue- and development-specific biology in sheep.

**Figure 2.**
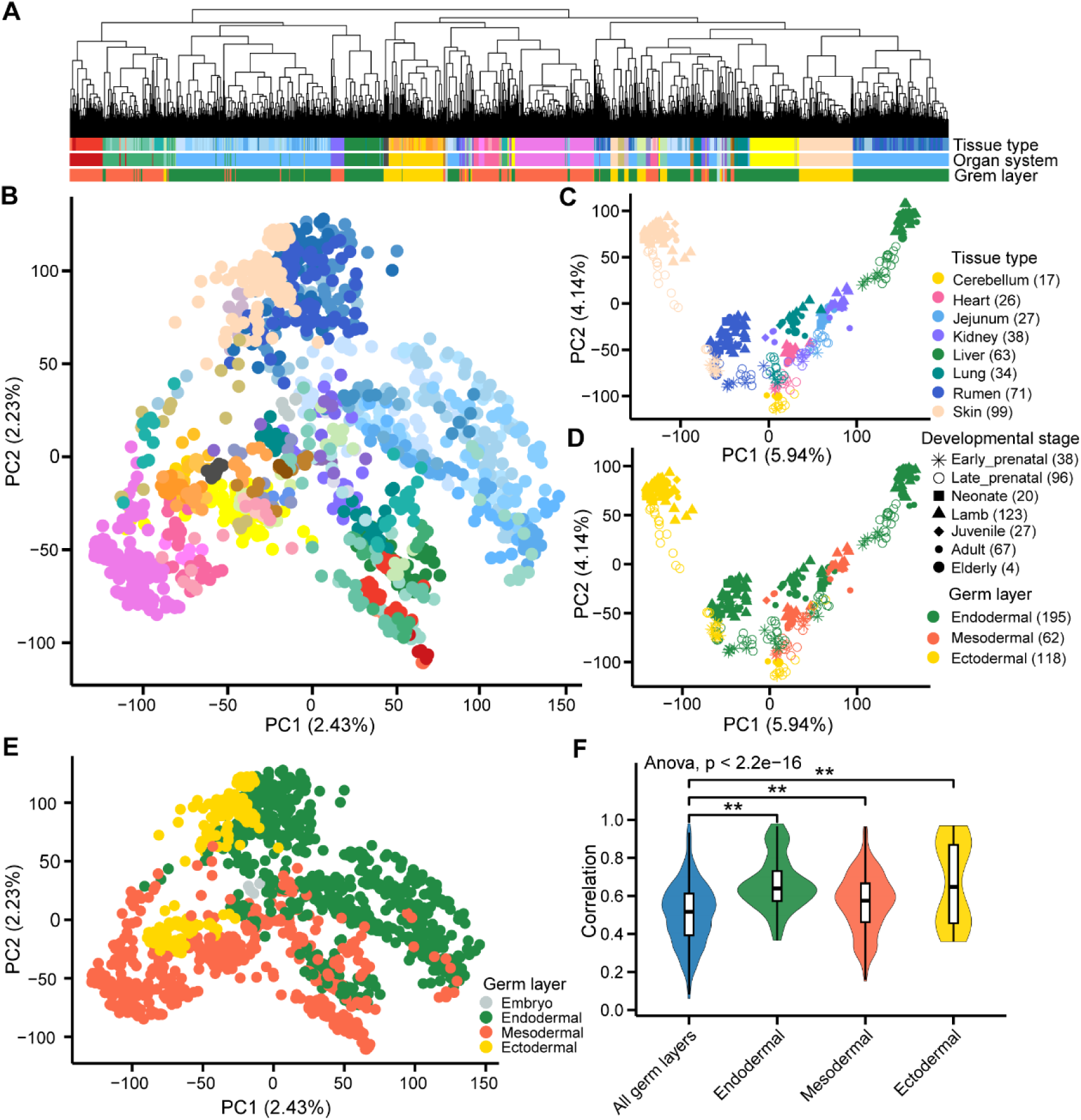
Clustering analysis of RNA-seq samples reveals tissue- and development-specific gene expression patterns. **A.** Hierarchical clustering of 1,413 RNA-seq samples based on distances (1-*r*, *r* is the Pearson correlation coefficient calculated from gene expression values [Transcripts per Million, TPM]) of 6,000 genes with the highest expression variance across samples (measured by the standard deviation of TPM). **B.** Principal component analysis (PCA) was performed on 1,413 RNA-seq samples using the scaled expression (i.e., log_2_(TPM+0.25)) of the same 6,000 genes above. **C.** and **D.** PCA of 375 samples of eight tissues derived from three germ layers, spanning all the seven developmental stages, i.e., early prenatal (≤ E70), late prenatal (>E70), neonatal (D0-D8), lamb (D8-M6), juvenile (M6-Y1.5) adult (Y1.5-Y7), and elderly (>7Y). The samples in the top and bottom panels were colored to represent tissue type and germ layer, respectively. The shapes represent developmental stages. **E.** PCA of 51 tissues based on the median gene expression of the same 6,000 genes above, representing embryo, endodermal, mesodermal and ectodermal lineages. **F.** Comparison of gene expression correlation between tissues from the same and all germ layers.

### Discovery of tissue-specific gene expression and alternative splicing patterns

By comparing a tissue with the rest, we considered the top 5% of upregulated genes in the target tissue based on FDR values as tissue-specific genes in each of 51 tissues (Figure 3A). In general, the functional enrichment analysis of these tissue-specific genes revealed known biological and physiological functions of the corresponding tissues (Figure 3B, Figure S6). For instance, adipose-specific genes were significantly (FDR < 0.05) enriched for cellular lipid catabolic process and fatty acid metabolic process, while cerebellum-specific genes were associated with nervous system development and neuron differentiation (Figure 3B). We also detected genes with tissue-specific alternative splicing patterns (Figure S7A), the function of which also reflected the known tissue biology (Figure S7B). For instance, 60 genes with specific alternative splicing in the cerebellum were significantly enriched in nervous system development. We further performed a motif enrichment analysis for promoters of these tissue-specific genes to detect the key transcription factors (TFs) determining tissue identity and functions. In general, the enriched motifs and expression of their target TFs revealed tissue-specific biology (Figure 3C). For example, the motifs of GATA6 TF were significantly enriched in cardiac muscle and ventricle-specific genes, and GATA6 also showed a specific expression in cardiac muscle and ventricle. GATA6 has been proposed to play an important role in heart development[33]. Similar findings were observed for ARNT2 in the hypothalamus and FOXF1 in the lung, which were also supported by previous evidence [34] (Figure 3C). Some novel findings included PROX1 in the liver, TBX3 in the placenta, and PATZ1 in the oviduct.

**Figure 3.**
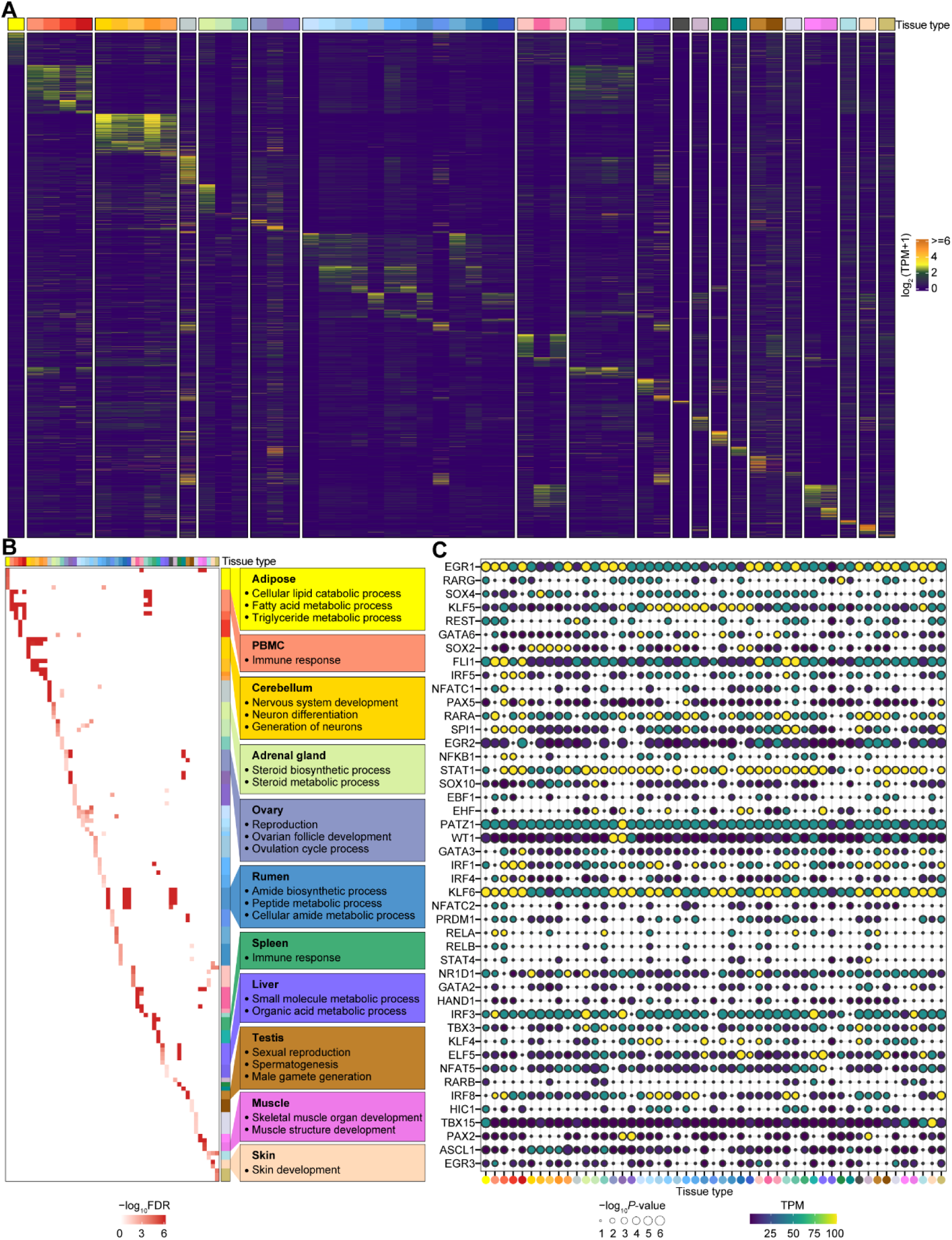
Tissue-specific gene expression. **A.** Inter-individual median expression levels of tissue-specific genes across 51 tissues. Each row represents a gene, and each column represents an individual tissue. Color represents the median log_2_(TPM+1) of genes among all the samples in a tissue. **B.** Gene Ontology (GO) terms significantly enriched (False Discovery Rate, FDR < 0.05) in tissue-specific genes that were identified across 51 tissues, with examples of significant GO terms for several tissues presented in the left boxes. **C.** Motifs of transcriptional factors (TFs) that were significantly (*P*< 0.05) enriched in promoters of tissue-specific genes. The x-axis represents the 51 tissues, with the same tissue color as presented in Fig 1A.

### Features of stage-specific genes in multiple tissues

We identified an average of 3,150 developmental stage-specific genes across 20 tissues (over three biological replicates per developmental stage per tissue) (Figure S8). In general, prenatal tissues had a larger number of stage-specific genes than postnatal ones (Figure S8). Notably, in all examined tissues, PCGs were the predominant gene type among the stage-specific genes, followed by lncRNAs, indicating the crucial role of PCGs in orchestrating developmental processes (Figure 4, Figure S9-11). Developmental stage-specific genes were significantly enriched in biological processes relevant to the developmental, anatomical, and physiological features of respective tissues (Figure 4, Figure S9-11). For instance, stage-specific genes in the muscle at early prenatal, late-prenatal, neonate, lamb, juvenile, and adult stages were significantly enriched for amide and peptide biosynthetic processes, muscle development and differentiation, various immune processes, body morphogenesis, mesenchymal development and the regulation of fibroblast growth (Figure 4A). Additionally, we revealed key genes that play pivotal roles in these transitions, along with stage-specific TFs that regulate developmental processes for each tissue (Figure 4, Figure S9-11). For instance, E2F1, SOX4, PLAG1, ARNT2, and MYOG are crucial for the proliferation and differentiation of muscle cells during the embryonic stage[35–38]. In the neonatal stage, KLF15, SOX10, and ZIC3 serve as essential TFs for muscle development and the maintenance of the myogenic lineage[39–41]. EGR1, SPI1, and POU2F2 are significantly enriched during the lamb stage, responding to growth factors and immune signals[42, 43]. In juveniles, REL is associated with immune responses and also plays a critical role in muscle development and regeneration[44]. In adults, ELF5 and PRDM6 are vital for regulating muscle cell differentiation and influencing the self-renewal and differentiation of muscle stem cells (Figure 4A) [45].

**Figure 4.**
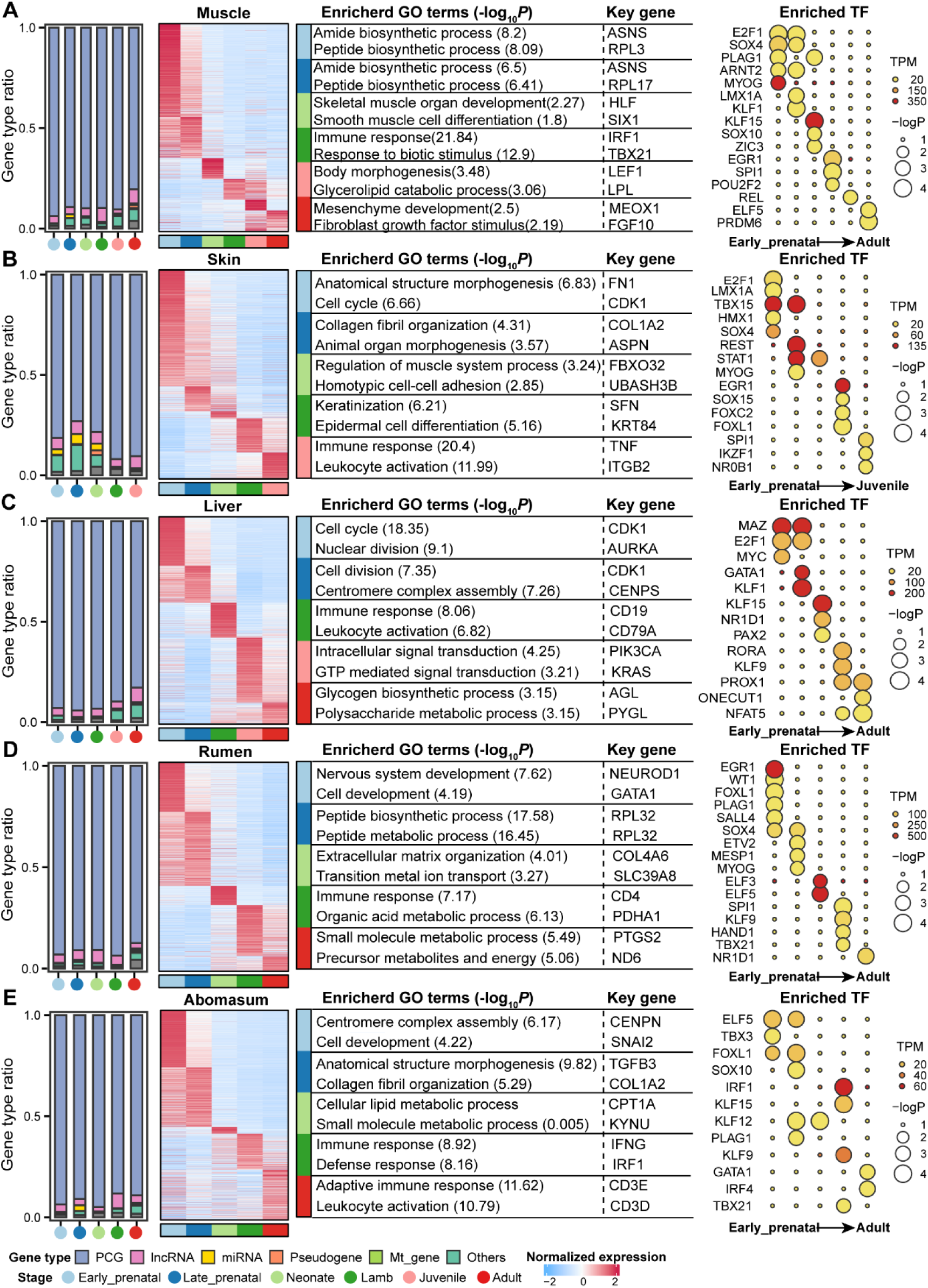
Features of stage-specific genes in major tissues. **A.** The panels for muscle from left to right illustrate the distribution of different gene types (protein-coding genes (PCGs), long non-coding RNAs (lncRNAs), microRNA (miRNAs), mitochondrial RNA (mtRNA), pseudogenes and others) among stage-specific genes; the expression level (log_2_(TPM+1)) of stage-specific genes across developmental stages; the biological processes enriched with stage-specific genes; motifs of transcriptional factors (TFs) that were significantly enriched in promoters of stage-specific PCGs, respectively. **B, C, D,** and **E** similar to **A** but for skin, liver, rumen, and abomasum, respectively. TPM: Transcripts per million.

### Gene clusters with distinct patterns of expression and TF regulation across developmental stages

To explore whether some genes share similar developmental expression patterns among the 20 tissues, we clustered genes according to their expression similarity across tissues using a soft-clustering approach (c-means), resulting in eight gene clusters (Figure 5A, Figure S12). For instance, 2,246 genes in cluster-2 exhibited a peak of expression at the lamb stage, and then showed a quick decrease at the later stages. Among them, 17.01% of genes were specifically and highly expressed in the skin, which was significantly enriched for cation transport and motifs of GATA3. GATA3 plays an important role in endothelial cell biology[46]. A total of 3,803 genes in cluster-7 were upregulated at the late-prenatal stage and then downregulated at the lamb stage, and 17.35% of them had a specifically high expression in the muscle. These genes were significantly enriched for the muscle system process and motifs of MYOD. MYOD participates in muscle cell differentiation[47]. The expression of 2,514 genes in cluster-8 increased continuously from the early-prenatal stage on, and 16.47% of them were specifically and highly expressed in the spleen and significantly enriched for cellular respiration and motifs of NFAT5. NFAT5 is engaged in inducible gene transcription during the immune response [48]. The expression of 3,708 genes in cluster-6 gradually increased as development progressed and upregulated sharply after the late-prenatal stage. Genes of this cluster were not dominated by tissue-specific genes of any tissues, which were significantly enriched with fundamental biological processes such as the small molecule metabolic process and motifs of SP1. SP1 is engaged in many cellular processes, including cell differentiation, cell growth, apoptosis, immune responses, and chromatin remodeling [49]. We also clustered genes using the same soft-clustering approaches within each of the 20 tissues (Figure 5B-E, Figure S13-15). We took rumen, muscle, skin, and liver as examples in Figure 5B-E. We identified seven, six, six and six gene clusters with distinct developmental expression patterns and enriched motifs in the rumen, skin, muscle, and liver, respectively (Figure 5B-E). For instance, the gene cluster-2 in the rumen encompassed 988 genes that displayed a gradual increase in expression as development progressed and were significantly enriched for processes related to the circadian rhythm and motifs of NR1D1 (Figure 5B). Beyond its well-known role in circadian rhythm regulation[50], *NR1D1* is also involved in several other biological processes, including metabolism[51], immune response[52], and cell differentiation[53], representing a high pleiotropic effect. In this study, we also found that NR1D1 might serve as a developmental stage-specific transcription factor (TF) across multiple tissues (Figure S16). However, future functional studies are required to confirm the role of *NR1D1* in developmental transitions. Of note, the expression pattern of these genes shared certain similarities between the rumen and skin but not muscle or liver across developmental stages, consistent with the notation that both rumen and skin are epithelial tissues (Figure 5F-I).

**Figure 5.**
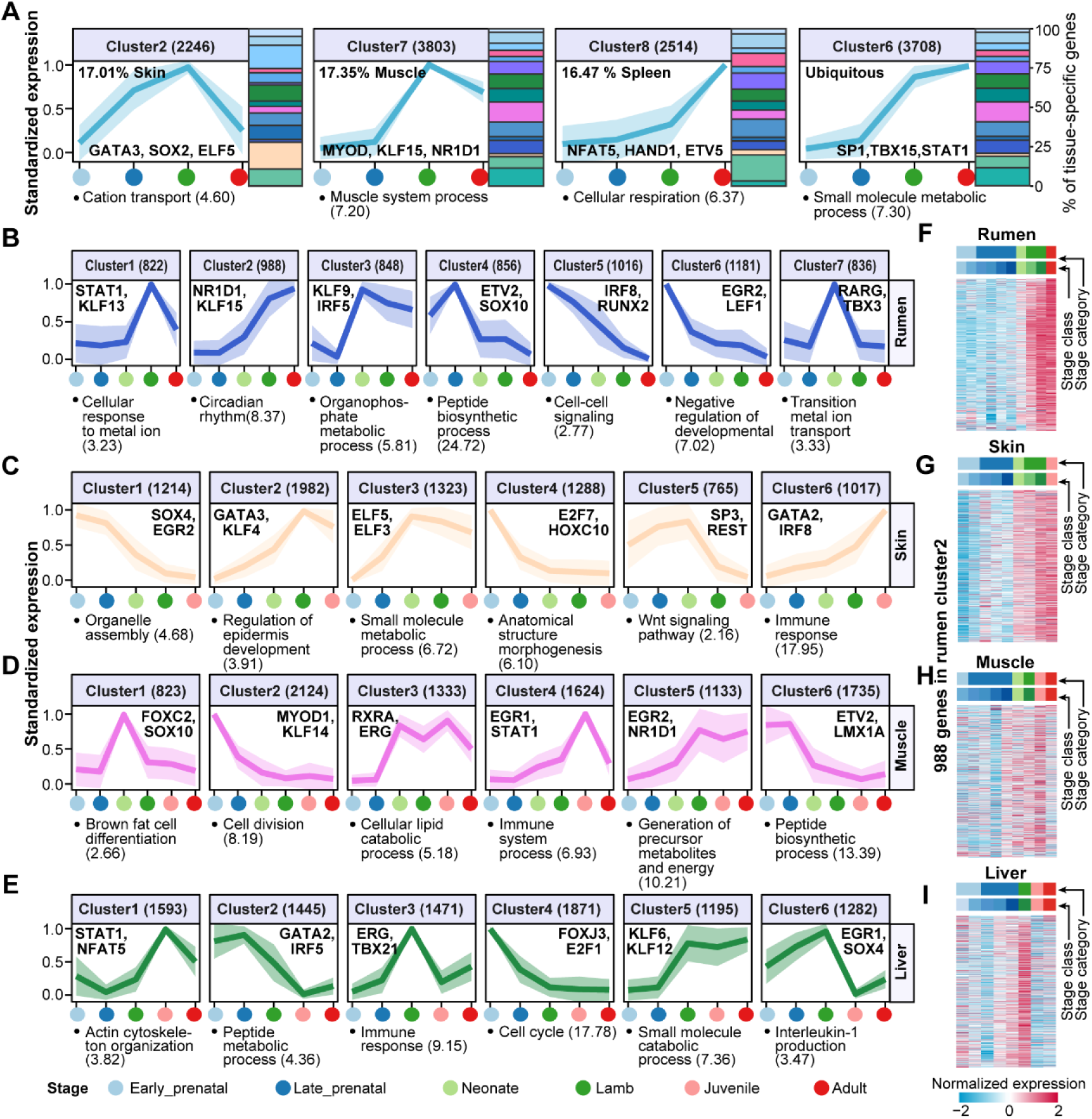
Time-course clustering analysis. **A.** Trajectory clustering of all tissue-specific genes in all 20 tissues across four developmental stages, i.e., early-prenatal, late-prenatal, lamb and adult stages. These trajectories were grouped into eight distinct gene clusters. Four clusters are displayed here as examples, and others are displayed in Figure S10. 17.01% of genes in cluster-2 are skin-specific, and 17.35% of genes in cluster-7 are muscle-specific, while 16.47% of genes in cluster-8 are spleen-specific. Genes in cluster 6 are not dominated by any tissues. The top significantly (FDR < 0.05) enriched transcription factors (TFs) of promoters of genes in each cluster are listed at the bottom of the respective panel. The y-axis represents the standardized expression values of tissue-specific genes in each of the clusters, while the x-axis represents developmental stages. The colors of developmental stages are shown at the bottom of the entire figure. The polygons represent mean ± standard deviation. Numbers in the bracket at the top of each panel indicate the number of genes in the respective cluster. The top significantly (FDR < 0.05) enriched Gene Ontology (GO) term for each cluster is listed at the bottom of the respective panel. **B.-E.** Similar to **A**, but for clustering of developmental stage-specific genes in the rumen, skin, muscle and liver, respectively. **F.-I.** Heatmaps show the normalized gene expression patterns of 998 genes detected in rumen cluster-2 across four tissues, i.e., rumen, skin, muscle, and liver, respectively.

### Gene co-expression analysis enhances functional annotation of the sheep genes

To improve the functional annotation of sheep genes using such a large and diverse transcriptome dataset, we have rephased this as below: We performed co-expression analyses using six complementary approaches in two scenarios i.e., 1) all the tissues together (integrated approach) and 2) each tissue separately (separated approach) [54–59]. In total, we identified 237 and 2,437 gene modules from the integrated approach and separated approach, respectively (Figure 6A, Figure S17A-C), and further evaluate the similarity and difference between the gene modules identified by different methods by calculating the Jaccard similarity coefficient (referred to here as the module sharing index), which is measured as the ratio of the intersection to the union of genes between module pairs (Figure S17D). We specifically compared the modules identified by WGCNA with those from the remaining methods (Figure S17E and F). In general, modules detected by these methods exhibited a low module sharing index, reflecting that they are complementary approaches for gene co-expression analyses[60, 61], as demonstrated by recent benchmarks that there was no single best co-expression analysis method. Among all the 2,437 modules detected by the single-tissue approach, 49.45% (1,205) were significantly enriched with at least one GO term (FDR < 0.05) (Figure 6B). Cross-tissue module preservation analysis identified 67 modules, of which 26 were tissue-shared and strongly preserved (Zsummary > 10), while 9 were tissue-specific with weak preservation (Zsummary < 2; Figure S18). Tissue-shared modules are more likely to be enriched with fundamental biological processes (e.g., cell cycle, chromatin assembly, translation), while tissue-specific modules tend to be enriched with tissue-relevant biology. For instance, two muscle-specific modules were significantly enriched in muscle development (*P*=7.98 × 10^−6^) and immune response (*P*=2.22 × 10^−43^) (Figure 6B). Among all the 26,423 genes in the 2,437 modules, 8,813 (33.35 %) genes had no functional annotation in the current Gene Ontology (GO) database (release 2022-01-18), which were referred to as “unannotated genes” hereafter. Compared to annotated genes, unannotated genes had lower expression and higher tissue-specificity, as well as were less likely to have human orthologous genes (Figure 6C). The proportion of unannotated genes in co-expression modules was distinct across tissues (Figure 6D). The embryo and mammary gland had the largest proportions of unannotated genes (an average of 33.15% and 31.14%, respectively). In contrast, central nervous system (CNS) and liver had the smallest proportions (an average of 18.70% and 19.01%, respectively). Of note, a total of 2,861 unannotated PCGs were able to be assigned to at least one co-expression module, contributing to their functional annotation (Figure S17G and H). For instance, in a module detected in muscle by WGCNA, 34 unannotated genes were co-expressed with 106 genes that were functionally annotated with muscle morphogenesis, skeletal muscle cell differentiation, and skeletal muscle development by the GO database (Figure 6E). We could thus infer that these 34 unannotated genes were more likely to have similar functions on muscle development in sheep.

**Figure 6.**
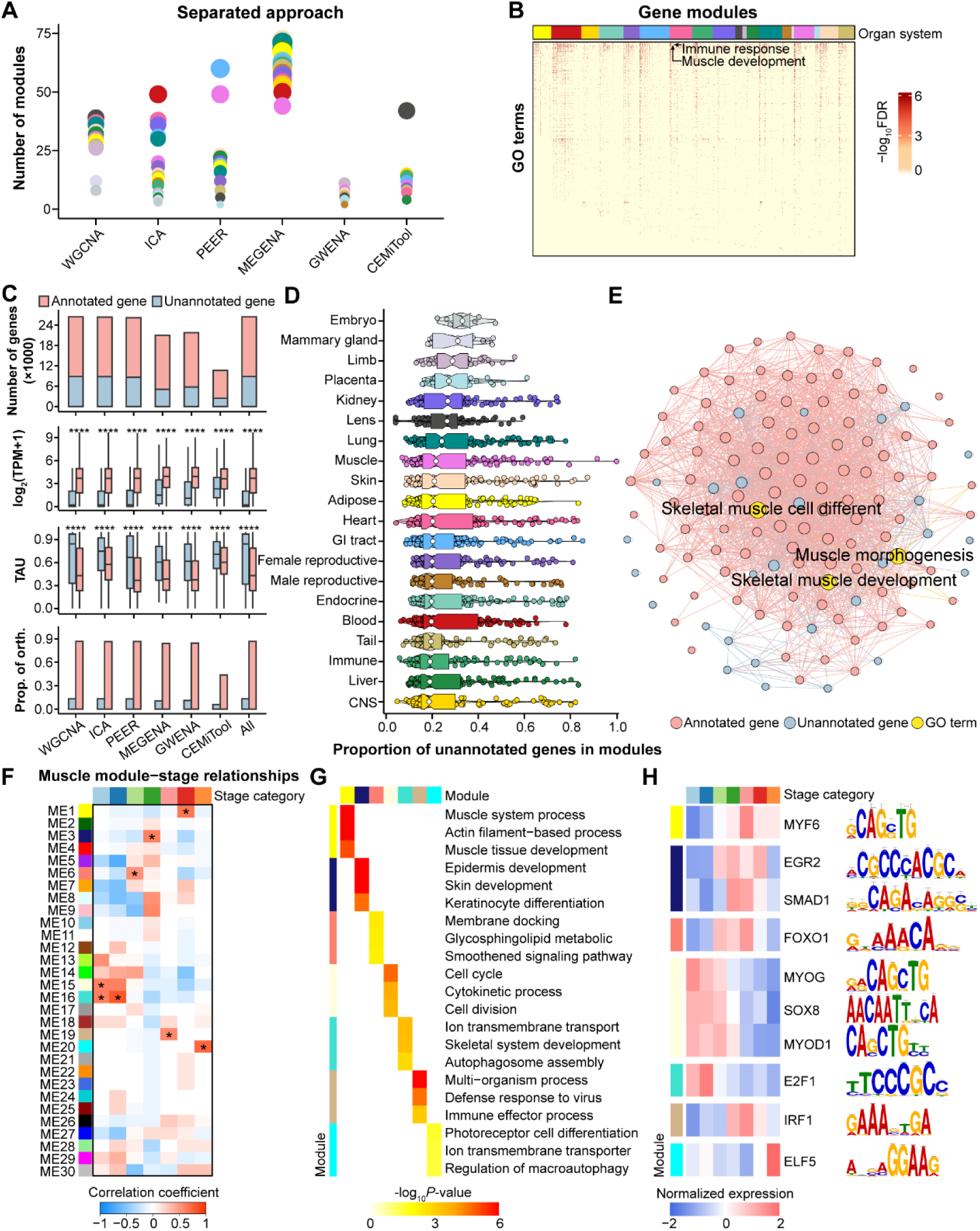
Gene co-expression network analysis across tissues and developmental stages. **A.** Number of gene modules detected in each of 20 tissues by six complementary gene co-expression approaches, including WGCNA, ICA, PEER, MEGENA, GWENA and CEMiTool. Each dot represents a tissue, and the dot size represents the number of detected gene modules. **B.** Significantly enriched GO terms (FDR < 0.05) in gene modules across 20 tissues. **C.** The comparison of genes with and without GO annotation (referred to as “annotated genes” and “unannotated genes”, respectively) in gene co-expression modules across different features. The gene functional annotation was based on the Gene Ontology database (version 2022-01-18). “All” means the combined results from the six co-expression methods. The plots from top to bottom are number of genes, expression level, TAU values, and the proportion of orthologous genes in humans. Significant differences between annotated and unannotated genes are obtained by a two-sided *t*-test. ** means *P* < 0.01. **D.** The proportion of unannotated genes in each of the gene co-expression modules across 20 tissues. **E.** An example of gene co-expression modules in muscle detected by WGCNA includes 34 unannotated and 106 annotated genes. All these genes are significantly (FDR < 0.05) enriched in muscle morphogenesis, skeletal muscle cell differentiation, and skeletal muscle development, which are denoted as yellow. The edges between genes represent Pearson’s correlations (r ≥ 0.7, *P* < 0.05) of expression levels across all the 129 samples in muscle. The circle sizes correspond to the interaction degrees. **F.** Correlations between gene modules and developmental stages in muscle. The statistical significance of the module-developmental stage relationship is corrected for multiple testing using the FDR method. The stars denote FDR < 0.05 (muscle-specific modules). **G.** The top significantly enriched GO terms (*P*<0.05) of developmental stage-specific modules in muscle. **H.** The top enriched motifs and expression patterns of transcriptional factors (TFs) of developmental stage-specific modules in muscle.

Within each of the 20 tissues, we further linked the WGCNA-detected co-expression modules to developmental stages. We took muscle as an example in Figure 6F-H. In total, seven of 30 co-expression modules were detected as developmental stage-specific modules (Figure 6F). For instance, genes in module-1 (ME1) were significantly (FDR < 0.05) upregulated at the adult stage and enriched in the muscle system process, actin filament-based process, and muscle tissue development (Figure 6G). The following motif enrichment analysis of genes in developmental stage-specific modules revealed biologically relevant transcription factors (TFs), which also showed developmental stage-specific expression (Figure 6H). For instance, MYF6, acting as a myogenic factor that prompts fibroblasts to differentiate into myoblasts[62], was observed with specific expression in adults. MYOG and MYOD1, well-recognized transcription factors essential for the development of functional skeletal muscle[63, 64], were highly expressed at the early-prenatal stage. Additionally, FOXO1, a key player in myogenic growth and differentiation[65, 66], demonstrated developmental stage-specific expression in neonates. Furthermore, the analysis proposed several novel TFs for muscle development for future functional validations. For instance, SOX8, involved in regulating embryonic development and determining cell fate, exhibited developmental stage-specific expression in muscle at the early-prenatal stage. In contrast, SMAD1, which plays a role in developmental processes and immune responses, showed developmental stage-specific expression in muscle at the lamb stage. Additionally, IRF1, functioning as an activator for genes implicated in innate and acquired immune responses, showed developmental stage-specific expression in muscle at the juvenile stage. Together, these findings advanced our understanding of the development-dependent roles of transcription factors in muscle development and function in sheep. In addition, we found that the STAT[67] and KLF[68] families played essential roles in regulating the development and function of the liver and kidney, respectively (Figure S19).

### Exploiting the dGEA to interpret rumen evolution and monogenic traits in sheep

Previous studies have identified a group of genes known as the epidermal differentiation complex (EDC), which are responsible for epithelial tissue (e.g., rumen, skin, and wool) development and repair by regulating the terminal differentiation program of the keratinocytes through a series of coordinated and inter-dependent signal transduction pathways [1, 69–71]. We examined expression patterns of all 18 EDC genes across tissues and developmental stages. Among them, *PGLYRP3*, *PGLYRP4*, *PRD-SPRRII*, *S100A12* and *S100A8* were specifically and highly expressed in the rumen, while *C1orf68*, *PRR9*, *S100A1*, *S100A3* and *TCHHL1* were specifically and highly expressed in the skin (Figure 7A). All five rumen-specific EDC genes showed similar developmental stage-specific expression patterns and the highest expression levels at postnatal day 7 (P7) (Figure 7B). All the five skin-specific genes have little or no expression at the early prenatal stages (i.e., embryo, E55–E85) and then rapidly upregulated after E85, partially because the number of primary follicles increases, and the secondary follicles start to form (Figure 7C).

**Figure 7.**
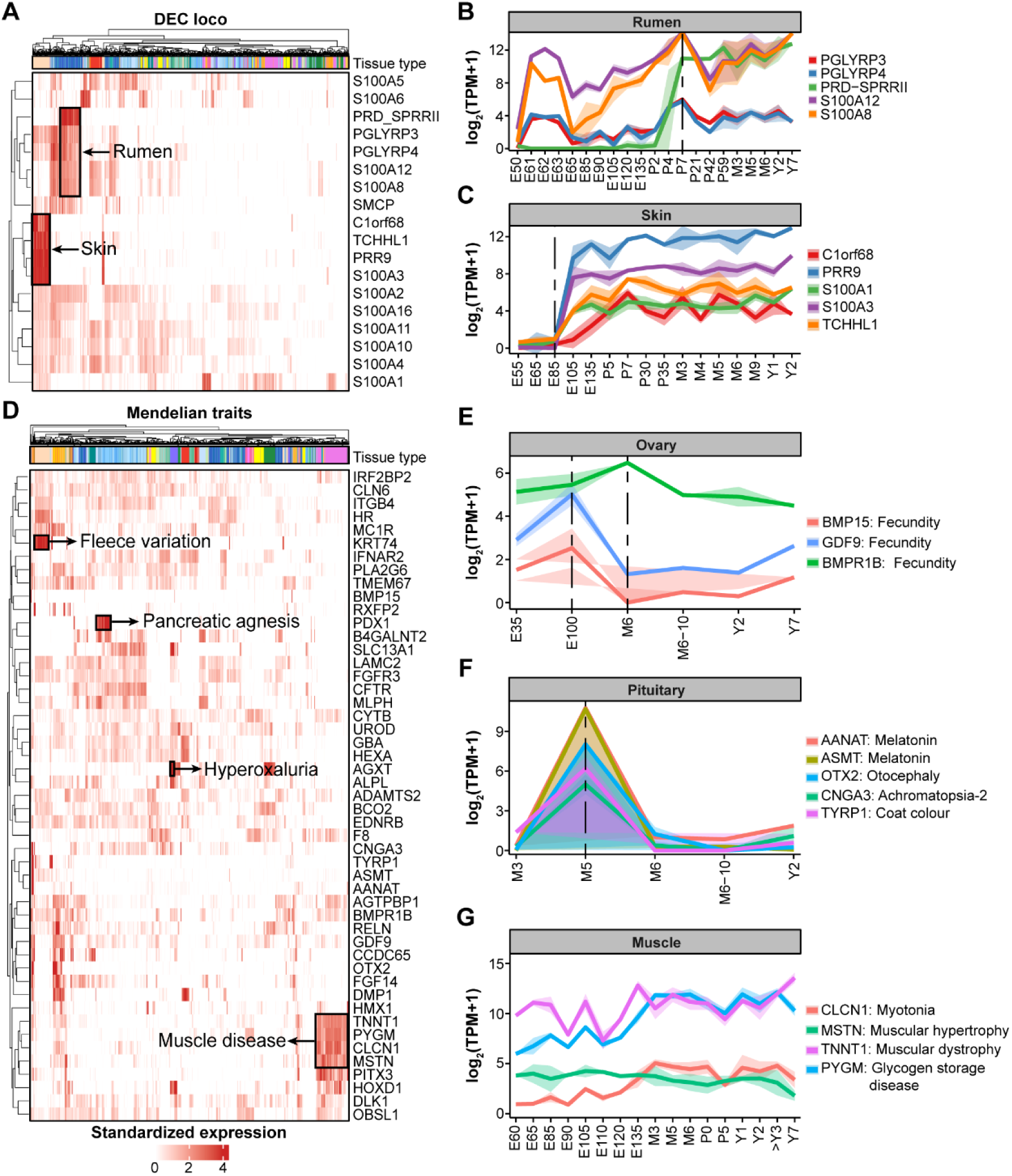
Exploiting the developmental Gene Expression Atlas (dGEA) to interpret ruminant-relevant genes and monogenic traits in sheep. Expression levels of 18 genes in the epidermal development complex (EDC) region across 51 tissues. The gene expression levels are normalized as transcripts per million (TPM). The y-axis represents the 18 EDC genes, and the x-axis represents the 51 tissues. **A.** The developmental expression patterns of five EDC genes with specific expression in rumen. **C** Similar to **B**, but another 5 EDC genes with specific expression in skin. **D.** Expression levels of 49 genes associated with sheep monogenic traits across the 51 tissues. **E.-G.** developmental expression patterns of genes with specific expression in ovary, pituitary, and muscle, respectively. The monogenic traits associated with each gene are listed next to it.

We examined the Online Mendelian Inheritance in Animals (OMIA)[72] to explore expression patterns of 49 causal genes of 48 monogenic traits across tissues and developmental stages in sheep. Many of them showed strong tissue and developmental stage-specific expression patterns (Figure 7D, Figure S20). For instance, *BMPR1B*, *BMP15*, and *GDF9* are proposed as causal genes of fecundity in sheep, which were specifically and highly expressed in the ovary (Figure 7D). However, the developmental expression patterns of *BMP15* and *GDF9* were different from that of *BMPR1B*. *BMP15* and *GDF9* showed the highest expression level at E100 and the lowest expression at six months ago (M6), whereas *BMPR1B* showed the highest expression level at M6 (Figure 7E). Of note, our analysis also proposed novel trait-tissue associations for future functional validations. For example, causal genes of Melatonin (*AANAT*), Melatonin (*ASMT*), Achromatopsia-2 (*CNGA3*), Otocephaly (*OTX2*) and Coat color (*TYRP1*) were specifically and highly expressed in the pituitary. These genes showed similar developmental patterns, with little expression at three months of age (M3) and a quick peak at M5 but then downregulated sharply afterwards, indicating their specific functional roles in the pituitary at M5 (Figure 7F). Similarly, five causal genes of muscle-relevant diseases, including *MSTN* and *DLK1* of muscular hypertrophy, *TNNT1* of muscular dystrophy, *CLCN1* of myotonia, and P*YGM* of glycogen storage disease, exhibited a strong muscle-specific expression, but showed different expression patterns as development progressed (Figure 7G). Furthermore, dGEA can also help interpret the tissue and developmental expression patterns of genes already annotated by GO terms in sheep. For instance, 12 genes engaged in the lipid metabolic process were explicitly and highly expressed in adipose (Figure S21A). The expression of these genes gradually increased from E85 and reached a peak at E132 before decreasing sharply, then rebounding by M5, and stabilizing at a high expression from that point onwards (Figure S21B).

### Genetic variants of complex traits were enriched in tissue- and developmental stage-specific genes

To explore whether the dGEA atlas can improve our understanding of the genetic and biological basis underlying complex traits in sheep, we first conducted single-marker GWAS for 12 complex traits of economic importance in a Merino sheep population (N =1,639), representing five wool traits, five reproductive traits and two growth traits. We then tested the enrichment of GWAS signals with tissue- and developmental stage-specific genes and gene clusters detected above (Figure 8, Table S2). As shown in Figure 8A, GWAS signals of complex traits were significantly (FDR < 0.05) enriched in genes with tissue-specific expression. In general, immune/blood-specific tissues showed the highest enrichment for all the traits, followed by gastrointestinal (GI) tract-specific genes. In contrast, CNS-specific genes showed no significant enrichment for any traits (Figure 8A). The muscle-specific genes showed significant enrichments for GWAS signals of mean staple length (MSL), gestation length (GL) and individual birthweight (IBW). Compared to other traits, GWAS signals of wool traits such as CV (coefficient of variation of the fibre diameter), CN (crimp number), MSL, and greasy fleece weight (GFW) were more likely to be enriched in Skin-specific genes. Of note, GWAS signals of all the complex traits showed higher enrichments in genes with specific expression in prenatal tissues than those in postnatal tissues (Figure 8B). For instance, genes with specific expression in the prenatal rather than adult rumen, lymph node, and skin were significantly associated with many traits (Figure 8C). Within the rumen, genes in cluster-1, the expression of which showed a peak at the lamb stage and quickly decreased from then on (Figure 8B), were significantly enriched with five traits, including CV, MSL, GL, total litter weight at birth (TLWB) and IBW (Figure 8D). While all the gene clusters except for cluster-5 were significantly associated with IBW, indicative of the complexity of body growth regulation (Figure 8D).

**Figure 8.**
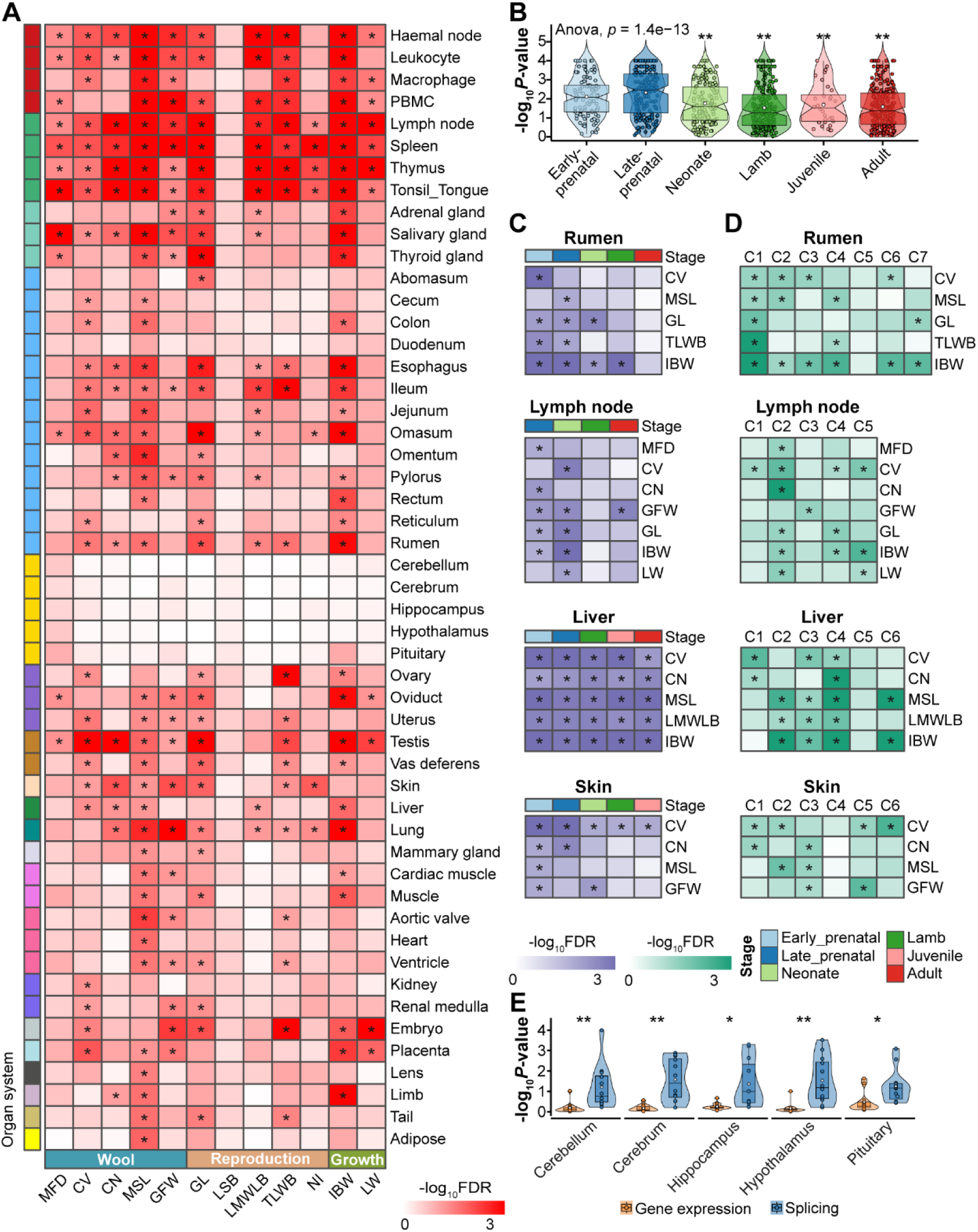
Enrichment of GWAS signals in genes with specific expression and splicing across various tissues and developmental stages. **A.** GWAS signal enrichment results of 12 complex traits in 51 tissues. The color corresponds to the enrichment degree (i.e., −log_10_FDR), which was computed by a sum-based GWAS signal enrichment analysis based on the tissue-specific genes, where (*) means FDR< 0.05. The x-axis represents the 12 complex traits i.e., MFD, mean fibre diameter; CV, coefficient of variation of the fibre diameter; CN, crimp number; MSL, mean staple length; GFW, greasy fleece weight; GL, gestation length; LSB, litter size at birth; LMWLB, litter mean weight per lamb born; TLWB, total litter weight at birth; NI, number of mating pregnancy; IBW, individual birthweight; LW, live weight. **B.** Comparison of GWAS signal enrichments (−log_10_*P*-value) of all the 12 complex traits in genes with developmental stage-specific gene expression. The ANOVA test was used to determine whether GWAS enrichments were significantly different across developmental stages, and the late-prenatal was selected as the target stage for pairwise comparisons with other stages. ** means *P* value < 0.01. **C** Similar to **A**, but for developmental stage-specific genes in rumen, lymph node, liver, and skin, respectively. **D** Similar to **C,** but for gene clusters associated with developmental stages. **E.** Comparison of GWAS signal enrichments (−log_10_*P*-value) between genes with tissue-specific expression and alternative splicing across cerebellum, cerebrum, hippocampus, hypothalamus, and pituitary. The paired *t*-test was used to determine whether GWAS signal enrichments were significantly different between these two groups of genes in each tissue, where (*) and (**) mean *P* value < 0.05 and 0.01, respectively.

Furthermore, we conducted GWAS signal enrichment analyses using alternative splicing results (Figure S22, Table S3). Like gene expression, genes with specific splicing patterns in many tissues were significantly enriched with GWAS signals of mean fibre diameter (MFD), MSL, TLWB, and IBW (Figure S22A). Of note, although genes with specific expression in CNS (cerebellum, cerebrum, hippocampus, hypothalamus, and pituitary) showed no enrichments for GWAS signals of any complex traits being studied (Figure 8A), genes with specific splicing patterns in CNS were significantly enriched for GWAS signals of MFD, CV, TLWB and IBW (Figure 8E). This finding proved the importance of studying alternative splicing of genes in CNS for understanding molecular mechanisms behind complex traits[73–75].

### Integrative analysis of genetic fine-mapping and sheep dGEA reveals key genes for complex traits in sheep

We conducted fine-mapping on QTL regions i.e., within ±500 kb of lead SNPs, to identify potential causal variants for sheep wool, growth, and reproductive traits (Table S4-5). In total, 196 SNPs were detected as causal variants (PIP > 0.9). By comprehensively integrating these fine-mapping results and the sheep dGEA, we identified several key genes associated with complex traits in sheep (Figure 9). For example, *SOX9* was associated with MSL and specifically expressed in skin. Its expression level gradually increased with developmental stages. *GNRHR* showed a strong association with LSB, specifically and highly expressed in the pituitary of adult sheep. *PRKDC* was associated with LW and showed the highest expression level at E135 in the spleen compared to other stages. In addition, we provided GO and KEGG functional enrichment for all the genes within ±500 kb of the identified causal variants (Table S6, Table S7).

**Figure 9.**
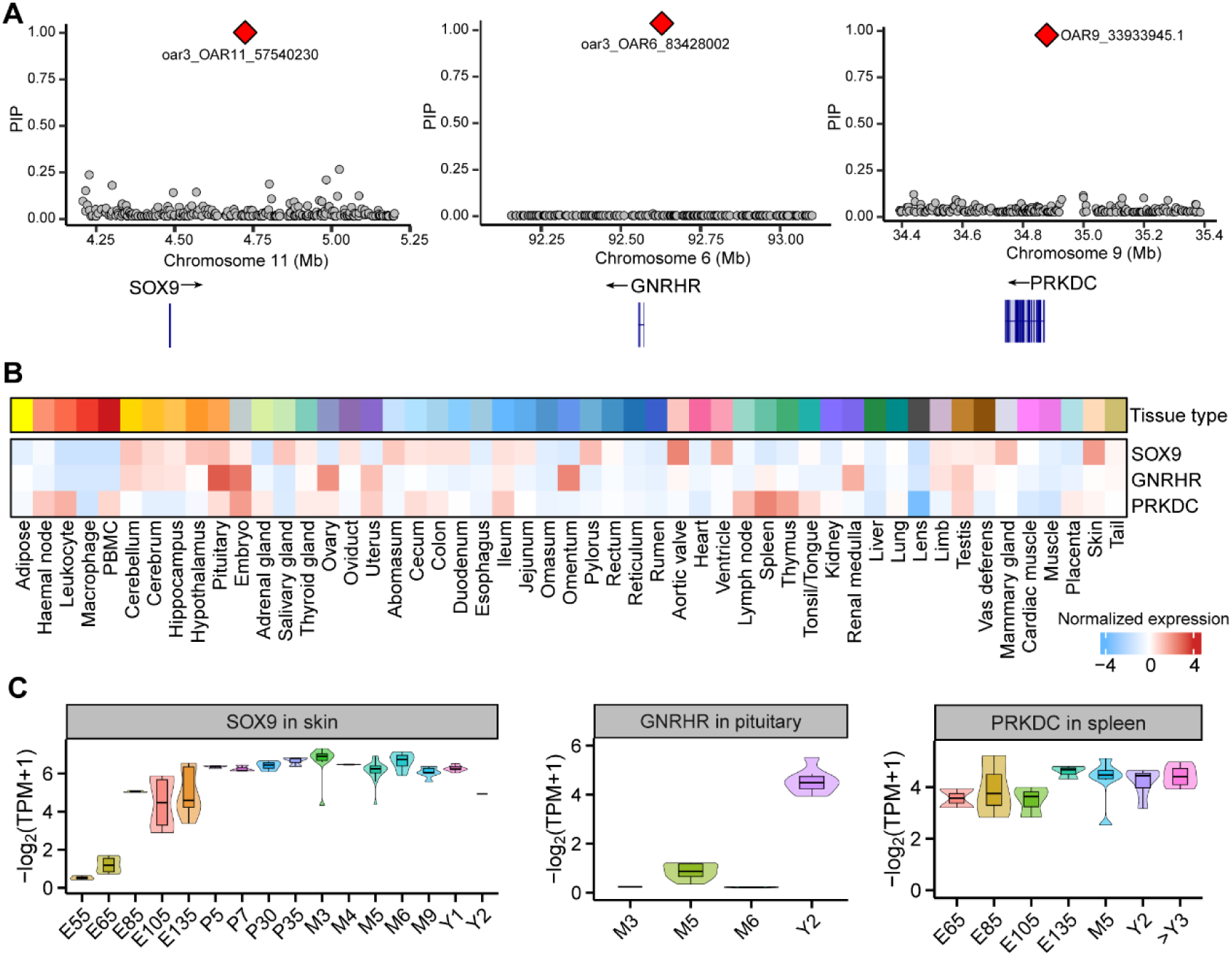
Integrative analysis of fine-mapping and sheep dGEA reveals key genes associated with complex traits in sheep. **A.** Fine-mapping for significant SNPs associated with wool, reproduction, and growth traits in sheep. **B.** Tissue-specific expression patterns of key genes. **C.** Stage-specific expression patterns of key genes.

## Discussion

In this study, we constructed a high-resolution multi-tissue developmental gene expression atlas in sheep by integrating 1,413 high-quality RNA-seq data, representing 51 tissues spanning 14 developmental time points from early organogenesis to adulthood, representing the largest study to date on multi-tissue developmental gene expression in sheep. We identified a series of developmentally dynamic PCGs that exhibited several signatures of functional enrichment. Our analysis revealed significant differences in the contribution of PCGs across different stages of organ development. Additionally, we found that complex traits in sheep are closely linked to the development of specific tissues. This extensive resource will greatly facilitate future research on more tissues and cell types in age-matched animals, as well as in disease challenge experiments, particularly in ruminants.

A fraction of protein-coding genes is house-keeping genes involved in fundamental biological processes, which are essential across many different tissues, such as cell cycle regulation, metabolism, and maintenance of cellular structure. In contrast, non-coding RNAs (ncRNAs) are frequently involved in more specialized regulatory roles, such as fine-tuning gene expression, often in a tissue-specific manner. Certain ncRNAs, particularly long non-coding RNAs (lncRNAs), have been found to exhibit high tissue specificity, as they may regulate gene expression in response to specific developmental signals or environmental stimuli in particular tissues[76]. Similar results have been reported in previous studies. For example, research by Jin et al. (2021)[76] demonstrated that lncRNAs tend to be more tissue-specific than protein-coding genes, which supports the observation that ncRNAs often play roles in fine-tuning tissue-specific gene expression. This has been observed across multiple organisms, including humans[77], cattle[20], pigs[21, 76], and chickens[78].

As described in previous studies [22, 32], we considered E70 as the point to distinguish the embryonic developmental stages. Given that the gestation period of sheep is approximately 150 days, E70 represents a midpoint in prenatal development. Additionally, around this point, significant transitions in organogenesis and physiological development occur, representing an important phase shift. Regarding the collection of tissues before E40, we acknowledge the challenges posed by their fragile nature and the difficulty in distinguishing anatomical structures at this early stage. Despite these complexities, we believe that the tissues collected prior to E40 still provide valuable information for early developmental processes, such as cell differentiation and the establishment of foundational organ systems. Although the sample size from stages prior to E40 is limited, we consider the data from these stages meaningful for downstream analyses, particularly for understanding early developmental events. Furthermore, we plan to collect more early-stage samples in future studies to enrich our dataset and provide a more comprehensive understanding of the developmental timeline.

Compared to the previous sheep gene expression atlas reported by Emily et al. in 2017 [22], we produced and analyzed approximately three times more RNA-seq data, covering nine more developmental stages, which enabled us to examine more genes (26,423 loci, among 20,471 protein-coding) than before (25,350 loci, among 19,921 protein coding). Moreover, while their study was limited to using network-based cluster analysis to group genes according to their expression pattern, we detected tissue and developmental stage-dependent gene expression, alternative splicing, and gene co-expression networks, and provided a significantly broader view of the dynamics of sheep gene expression. Furthermore, other studies have mainly focused on elucidating the genetic basis of high-altitude adaptation in sheep across 19 tissues[79] and 12 tissues[80], respectively. In contrast, we then, for the first time, exploited this extensive functional genome resource to decipher molecular and developmental mechanisms underlying both monogenetic and complex traits in sheep. We believe this valuable dGEA resource will significantly contribute to genetic and genomic research, selective breeding, and developmental biology in sheep.

By comprehensively integrating these fine-mapping results and the sheep dGEA, we identified several key genes associated with complex traits and provided novel underlying biological mechanisms in sheep. For instance, *SOX9*, a candidate gene for MSL in sheep, was mainly expressed in the skin. *SOX9* has been reported to be essential for outer root sheath differentiation and the formation of the hair stem cell compartment [81]. *GNRHR* was associated with LSB and showed a specifically high expression in the pituitary of adult sheep. Previous studies have demonstrated that the proper expression of gonadotropin-releasing hormone receptors (GnRHRs) by pituitary gonadotropes is critical for maintaining maximum reproductive capacity. In addition, GnRHR could influence the litter size of pigs [82] and goats[83]. Therefore, we proposed GnRHR as a strong candidate gene for litter size in sheep. A previous GWAS conducted by Horodyska et al. [84] revealed that *PRKDC* was associated with feed conversion efficiency and growth rate in pigs. It was also reported to be associated with body conformation traits in sheep [85]. We thus suggested that *PRKDC* could be a promising candidate gene for LW in sheep. In our study, gene expression patterns were generally conserved across breeds, as observed through clustering samples by tissue type and developmental stage rather than by breed, indicating that the phenotypic variation introduced by breed-specific gene expression is limited compared to tissue-specific and developmental-specific. In the future study, it is of interest to explore breed-specific gene expression and regulation by collecting large-scale healthy samples at the same tissue and developmental stages across distinct breeds.

We also noticed some limitations in the current study and give relevant perspectives below. Our initial assumption was that genomic variations ultimately influence complex traits by modifying the expression of genes in specific tissues and cell types with developmental stages. Previous research has demonstrated that the majority of expression quantitative trait loci (eQTL) are cis-variants, as reported by the GTEx Consortium [14]. Consequently, we concentrated on investigating cis-regulators of tissue-specific genes by expanding the analysis to include a specific range of distances (e.g., 20 kb) around these genes. To explore trans QTLs, however, a substantial number of samples is required for each tissue and cell type due to their relatively minor effects[9]. As tissues are a mixture of different cell types, the cell type composition of tissues being analyzed could confound our interpretation of some results. As we showed in Figure 8A, leukocyte, macrophage, and PBMC had distinct enrichments across wool, reproduction and growth traits. Therefore, it will be crucial to generate pure bulk cells and/or single-cell expression data to investigate which cell types across tissues and developmental stages are causal in a trait-relevant tissue. Furthermore, our ability to detect trait-relevant tissues and cell types is constrained by the limited availability of transcriptomic data. Consequently, we may inadvertently overlook tissues and cell types that are biologically significant for the traits of interest during specific physiological stages or under certain environmental conditions.

In the future, integrating more advanced technologies, such as single-cell omics[86–88], spatial transcriptomics[89, 90], long-read RNA-seq[91] or epigenomics[92], will further refine the functional annotation of sheep genes. In turn, this will help us further understand mammalian development and determine the causal molecular factors that drive organ development and influence organismal phenotypes[31, 93]. We hope to gain access to a greater variety of molecular phenotypes across different tissues and developmental stages of livestock, such as those being studied in the ongoing FarmGTEx project[20, 21]. By utilizing our current research strategy, we will be able to uncover new and valuable information about the genetic and biological factors that influence important agricultural traits. These efforts will significantly expand our knowledge of the genetic architecture and developmental basis underlying phenotypes in economically important mammalian species such as sheep.

## Materials and methods

### Tissue sample collection

In this study, we totally collected 426 tissue samples from two populations, including Chinese Merino and Australian Merino. The newly generated RNA-seq dataset from the Chinese Merino sheep consisted of 277 samples. The 12 embryos were collected from pregnant ewes at four embryonic time points (i.e., Day-65 embryos (E65), day-85 embryos (E85), day-105 embryos (E105), and day-135 embryos (E135); three biological replicates per time point). Specifically, we obtained 41 samples from E65 across 17 tissues, 78 samples from E85 across 29 tissues, 79 samples from E105 across 29 tissues, and 79 samples from E135 across 29 tissues.

The newly generated RNA-seq dataset comprised 149 samples for the Australian Merino sheep. Tissue samples were collected at eight different developmental stages, including four embryonic and four postnatal time points. The 65 embryonic samples included five day-30 (E30) embryos for esophageal tube, five day-50 embryos (E50) for rumen, five day-90 embryos (E90) for four tissues (reticulum, rumen, omasum, and abomasum), and five day-120 embryos (E120) for seven gut tissues (reticulum, rumen, omasum, abomasum, duodenum, cecum, and colon). The 84 postnatal samples consisted of the same seven gut tissues, collected from three neonatal lambs (before their first feed), three lambs at three weeks of age (before rumination, with no grass present in their GI tract), three at six weeks of age (once rumination established), and three at 12 weeks of age (after weaning). All tissues were collected immediately after the euthanasia of the ewes and stored in liquid nitrogen. The total RNA was extracted using the TRIzol reagent (Invitrogen), and sequencing libraries were prepared with the NEBNext Ultra RNA Library Prep Kit of Illumina (New England Biosystems). The RNA libraries were sequenced to generate 150-bp paired-end (PE150) reads using the Illumina NovaSeq 6000.

Furthermore, we downloaded 1,131 existing RNA-seq data (fastq files) within detailed metadata of developmental stages and tissue types from the NCBI Sequence Read Archive (SRA, https://www.ncbi.nlm.nih.gov/sra). In total, we analyzed 1,557 RNA-seq samples in this study. Based on known biology[9], we classified 51 tissues and cell types into 20 tissue categories. Details of all the RNA-seq samples are summarized in Table S1.

### Quantification of gene expression and alternative splicing

We analyzed all the 1,557 RNA-seq samples using the following bioinformatics pipeline. First, we removed adaptors and trimmed low-quality reads using Trimmomatic (v0.39) with the parameters of “TruSeq3-PE.fa:2:30:10 LEADING:3 TRAILING:3 SLIDINGWINDOW:4:15 MINLEN:36”[94]. We then aligned the clean reads to the sheep (Ovis aries) reference genome (Oar_rambouillet_v1.0, Ensembl version 104) using STAR (v2.7.8a)[95] with the parameters of “--quantMode GeneCounts --chimSegmentMin 10 --chimOutType Junctions --chimOutJunctionFormat 1 --outSAMtype BAM SortedByCoordinate --outSAMunmapped Within --readFilesCommand zcat --outFilterMismatchNmax 3”, resulting in an averaged uniquely mapping rate of 81.79% (Supplemental Table S1). For downstream analyses, we kept 1,470 samples with a uniquely mapping ratio >= 0.6 and clean reads count >10 million. We extracted raw read counts of 26,478 Ensembl (v104) genes by featureCounts (v2.0.1)[96] and obtained their normalized expression (i.e., TPM) using Stringtie (v2.1.4)[97], Salmon (v 1.3.0)[98] and Kallisto (v 0.45.1)[99]. We removed 6 samples with less than 40% of all the genes expressed (TPM ≥ 0.1), and three tissues with sample size < 3, resulting in 1,461 samples. We filtered genes that were not expressed in any samples, resulting in 26,423 genes for subsequent analyses. We performed the hierarchical clustering of all the RNA-seq samples using the *hclust* function in R package (v4.2.2). The distance between samples was measured by 1-*r*, where *r* was Pearson’s correlation coefficient based on the normalized quantile log_2_(TPM+0.25) of 6,000 genes with the highest variance across samples. We also visualized these samples using the PCA approach implemented in the *prcomp* function in R package. After filtering out 48 outliers based on sample clustering, we eventually kept 1,413 samples for subsequent analysis. For alternative splicing, we applied Leafcutter (v0.2.9)[100] to quantify intron excision levels using exon-exon junction reads from RNA-seq data in each of the tissue types as described previously[21]. We normalized intron excision levels as percent splicing (PSI) values for the downstream differential splicing (DS) analysis *via* the script “leafcutter_ds.R” in the LeafCutter package.

### Tissue-specific analysis of gene expression

We employed tspex to quantify the tissue specificity of gene expression by calculating TAU (τ) values [101]. By comparing a tissue with the remaining, we conducted gene differential expression analysis using the R package edgeR (v 3.40.2)[102], and considered the top 5% of upregulated genes in the target tissue based on FDR values as its tissue-specific genes [9]. We then conducted the functional enrichment analysis of tissue-specific genes for biological process (BP) terms in the gene ontology (GO) database using both R package clusterProfler [103]and gene set enrichment analysis (GSEA) [104], and considered FDR < 0.05 as significant. We performed the motif enrichment analysis for promoter regions (i.e., 1500 bp upstream and 500 bp downstream of transcriptional start sites, TSS) of tissue-specific genes using MEME software[105], based on the JASPAR (2020) core non-redundant vertebrate motifs[106] and the HOCOMOCO v11 human motif set[107].We consider FDR<0.05 as a significant motif.

### Development-specific and time-course gene expression analysis

To identify genes with developmental stage-specific expression, we carried out differential expression analysis between one stage and the rest in each of the 20 tissues using the R package edgeR (v 3.40.2) [102], where the sample size of each stage pre-tissue was > 3. We considered genes with log_2_FC > 2 and FDR < 0.05 as developmental stage-specific genes. Within each of the 20 tissues, we then clustered genes based on their expression levels across developmental stages using the soft-clustering approach (c-means) implemented in the R package mFuzz (v.2.32.0)[108, 109]. For this analysis, we only considered genes with development-specific expression detected above. The number of gene clusters was individually determined in each of 20 tissues using the minimum centroid distance measurement[108, 109].

### Gene co-expression network analysis

We applied six complementary methods with default parameters, including WGCNA (v1.69)[54], GWENA (v1.8.0)[55], ICA (v1.0.2)[56], PEER (v1.3)[57], MEGENA (v1.3.7)[58], and CEMiTool (v1.8.3)[59], to identify modules of co-expressed genes based on the quantile normalized gene expression. Gene co-expression clustering methods, such as WGCNA, GWENA and CEMiTool, aimed to identify non-overlapping co-expressed gene modules. In contrast, matrix factorization methods, such as ICA, PEER and MEGENA, used factor loadings as module eigengene profiles, resulting in larger modules that included overlapping sets of genes. Based on the background dataset of 26,423 genes, we further removed genes with zero standard deviation in each tissue to ensure the robustness of the analysis. Functional enrichment analysis of gene co-expression modules was conducted by using clusterProfiler (v4.0)[103], and the following visualization was done using Gephi (v0.9.2)[110]. To evaluate the similarity and difference between the gene modules identified by these methods, we calculated the Jaccard similarity coefficient (referred to here as the module share index)[111], which is measured as the ratio of the intersection to the union of genes between all module pairs.

### Cross-tissue module preservation analysis

To evaluate the preservation of cross-tissue modules in the network, we used the modulePreservation function in the WGCNA R package[112]. Zsummary composite preservation scores were calculated by averaging the several preservation statistics generated through many permutations of the raw data, which summarized the evidence that a module is preserved and indicative of module robustness and reproducibility[112]. Generally, modules with Zsummary scores >10 are considered strongly preserved (that is, tissue-shared modules), Zsummary scores between 2 and 10 are weakly to moderately preserved, and Zsummary scores <2 are not preserved (tissue-specific modules)[112, 113].

### GWAS and fine mapping

We performed the single-marker GWAS for five wool traits (MFD, mean fibre diameter; CV, coefficient of variation of the fibre diameter; CN, crimp number; MSL, mean staple length; GFW, greasy fleece weight), five reproduction traits (GL, gestation length; LSB, litter size at birth; LMWLB, litter mean weight per lamb born; TLWB, total litter weight at birth; NI, number of mating pregnancy) and two growth traits (LW, live weight; IBW, individual birthweight) in a population of 1,639 sheep with genotypes of the 600K arrays (n=504,805 SNPs). For wool traits and LW, we used the adjusted phenotypes (regressing out fixed effects, including flock, birth-year, and month) for GWAS analysis [7]. For reproduction traits and IBW, we used de-regressed proof values (DRP, breeding values) as phenotypes. Genotype imputation was performed using the BEAGLE (v5.1) software [114], and we only considered SNPs with call rate > 90%, minor allele frequency (MAF) >1%, *P* _(Hardy–Weinberg equilibrium)_ > 10^−6^ [7]. We then employed the linear mixed model, implemented in GCTA (v1.94.1)[115] software to test for the association of genomic variants with all the complex traits. For each trait, we considered the false discovery rate (FDR) < 0.05 as a threshold to identify the significant SNPs[116].

Plink software was employed to construct the LD matrix for the variants within a 500kb window flanking each significant SNP (identified in the above GWAS analysis)[117]. Subsequently, the susieR R package (version 0.12.35) was utilized to identify the causal variants linked to the significant SNPs by analyzing the summary statistics of GWAS through the susie_rss function[118]. The causal effect (as indicated by a posterior inclusion probability, PIP) was computed following this. Locus plots with gene annotations were created using the locuszoomr R package (version 0.3.5)

### Enrichment analysis of GWAS signals in tissue and development-specific genes

We applied the sum-based marker-set test approach below, as implemented in the R package QGG [119] to identify whether the GWAS signals were significantly enriched in a gene set, including tissue-specific genes, development-specific genes and co-expression modules:

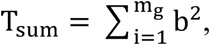

where T_sum_ is the summary statistics of a gene set, mg is the number of SNPs within the gene set, and b is the SNP effect from single-marker GWAS. We considered SNPs within 20kb up and downstream of the gene body to cover the *cis*-regulatory regions of genes. To obtain an empirical *P*-value for the association of a gene set with a complex trait, we employed a 10,000 times permutation strategy and then applied a one-tailed test of the proportion of random summary statistics greater than that observed, as described previously [9, 28]. We then corrected *P-*values for multiple testing using the FDR method and considered FDR < 0.05 significant.

### Ethical statement

All animal experiments were carried out following the ARRIVE Guidelines and the Regulations for the Administration of Affairs Concerning Experimental Animals (revised in March 2017) approved by the State Council of the People’s Republic of China, and also approved by the Animal Ethical and Welfare Committee of Xinjiang Academy of Animal Sciences (Approval No. 2019009). The Australian animal experiment was conducted at the University of New England, Armidale, Australia, in accordance with the University of New England Animal Ethics Committee approval (AEC no14-041)[120]. The pregnant ewes were housed indoors for a 7-day “settling in period” prior and allowed access to feed and water *ad libitum* and were not fed the night before they were slaughtered to being euthanized (electrocution and exsanguination).

## Data availability

All the 426 newly generated raw data in this study have been deposited in the NCBI SRA database under accession code PRJNA1006307. All raw data analyzed in this study are publicly available for download without restrictions from SRA (https://www.ncbi.nlm.nih.gov/sra/) under accession codes PRJEB19199, PRJEB6169, PRJNA485657, PRJNA414087, PRJNA287258, PRJNA309284, PRJNA315011, PRJNA362606, PRJNA395156, PRJNA432669, PRJNA450309, PRJNA490799, PRJNA504353, PRJNA524671, PRJNA526287, PRJNA596252, PRJNA638028, PRJNA665351, PRJNA729910, PRJNA736945, PRJNA182914, PRJNA244152, PRJNA378408, PRJNA421633, PRJNA507468, PRJNA565839, PRJNA595784 PRJNA625152, PRJNA639853, PRJNA705554. Details of RNA-seq datasets can be found in Table S1. The metadata and expression estimates relating to all the RNA-seq samples have been deposited on the Figshare website (10.6084/m9.figshare.23944452). Each use of software programs has been clearly indicated and information on the options that were used is provided in the Methods section.

## CRediT authorship contribution statement

**Bingru Zhao:** Methodology, Validation, Formal analysis, Investigation, Writing – original draft, Writing – review & editing. **Hanpeng Luo:** Software, Formal analysis, Data curation. **Xuefeng Fu:** Validation, Investigation, Resources. **Guoming Zhang:** Visualization, Investigation. **Emily L. Clark:** Data curation, Writing – original draft & editing. **Feng Wang:** Validation, Investigation. **Brian Paul Dalrymple:** Data curation, Writing – review & editing. **V. Hutton Oddy:** Data curation, Writing – review & editing. **Philip E. Vercoe:** Data curation, Writing – review & editing. **Cuiling Wu:** Resources. **George E. Liu:** Resources. **Cong-jun Li:** Resources. **Ruidong Xiang:** Formal analysis, Writing – review & editing, Funding acquisition. **Kechuan Tian:** Conceptualization, Writing – review & editing, Funding acquisition, Project administration. **Yanli Zhang:** Conceptualization, Formal analysis, Writing – review & editing, Funding acquisition, Project administration. **Lingzhao Fang:** Conceptualization, Methodology, Formal analysis, Investigation, Writing – review & editing, Project administration. All authors have read and approved the final manuscript.

## Acknowledgements

This work was supported by grants from the National Key Research and Development Program (No. 2021YFF1000702), the National Natural Science Foundation of China (No. 32402755), the Project of Seed Industry Revitalization in Jiangsu Province (No. JBGS[2021]113), the Technological Innovation Project of Shandong Academy of Agricultural Sciences (No. CXGC2023F10), the 2023 Excellent Postdoctoral Program in Jiangsu Province, China (No. 2023ZB584), and the high-performance computing platform of Bioinformatics Center, Nanjing Agricultural University. Additional support came from the Department of Agriculture, Fisheries and Forestry of the Commonwealth of Australia, Filling the Research Gap (2013-2016) DAFF 1193857-31 ‘Host Control of Methane Emissions from Sheep’, the Australian Research Council’s Discovery Projects (DP160101056 and DP200100499) supported by R.X, the Institute Strategic Programme grants BBS/E/D/10002070 and BB/X010945/1 awarded to the Roslin Institute by the Biotechnology and Biological Sciences Research Council supported by E.L.C. We would like to express our deep gratitude to the late Professor Shengli Zhang for his invaluable support and guidance.

## References

1. Jiang Y, Xie M, Chen WB, Talbot R, Maddox JF, Faraut T, et al. The sheep genome illuminates biology of the rumen and lipid metabolism. Science. 2014;344:1168–73.

2. Meuwissen T, Hayes B, Goddard M. Genomic selection: a paradigm shift in animal breeding. Animal frontiers. 2016;6:6–14.

3. Kalds P, Huang S, Xi S, Fang Y, Gao Y, Zhou S, et al. ABE-induced PDGFD start codon silencing unveils new insights into the genetic architecture of sheep fat tails. J Genet Genomics. 2023;8527:1.

4. Li X, He SG, Li WR, Luo LY, Yan Z, Mo DX, et al. Genomic analyses of wild argali, domestic sheep, and their hybrids provide insights into chromosome evolution, phenotypic variation, and germplasm innovation. Genome Research. 2022;32:1669–84.

5. Li X, Yang J, Shen M, Xie XL, Liu GJ, Xu YX, et al. Whole-genome resequencing of wild and domestic sheep identifies genes associated with morphological and agronomic traits. Nature Communications. 2020;11:2815.

6. Zhu HZ, Wu ZY, Ding X, Post MJ, Guo RP, Wang J, et al. Production of cultured meat from pig muscle stem cells. Biomaterials. 2022;287:121650.

7. Zhao BR, Luo HP, Huang XX, Wei C, Di J, Tian YZ, et al. Integration of a single-step genome-wide association study with a multi-tissue transcriptome analysis provides novel insights into the genetic basis of wool and weight traits in sheep. Genetics Selection Evolution. 2021;53:56.

8. Hu ZL, Park CA, Reecy JM. Bringing the animal QTLdb and CorrDB into the future: meeting new challenges and providing updated services. Nucleic Acids Research. 2022;50:D956–D61.

9. Fang LZ, Cai WT, Liu SL, Canela-Xandri O, Gao YH, Jiang JC, et al. Comprehensive analyses of 723 transcriptomes enhance genetic and biological interpretations for complex traits in cattle. Genome Res. 2020;30:790–801.

10. Finucane HK, Reshef YA, Anttila V, Slowikowski K, Gusev A, Byrnes A, et al. Heritability enrichment of specifically expressed genes identifies disease-relevant tissues and cell types. Nature Genetics. 2018;50:621–29.

11. Birney E, Stamatoyannopoulos JA, Dutta A, Guigo R, Gingeras TR, Margulies EH, et al. Identification and analysis of functional elements in 1% of the human genome by the ENCODE pilot project. Nature. 2007;447:799–816.

12. Ardlie KG, DeLuca DS, Segre AV, Sullivan TJ, Young TR, Gelfand ET, et al. The genotype-tissue expression (GTEx) pilot analysis: multitissue gene regulation in humans. Science. 2015;348:648–60.

13. Mele M, Ferreira PG, Reverter F, DeLuca DS, Monlong J, Sammeth M, et al. The human transcriptome across tissues and individuals. Science. 2015;348:660–65.

14. Aguet F, Brown AA, Castel SE, Davis JR, He Y, Jo B, et al. Genetic effects on gene expression across human tissues. Nature. 2017;550:204–13.

15. Aguet F, Barbeira AN, Bonazzola R, Brown A, Castel SE, Jo B, et al. The GTEx Consortium atlas of genetic regulatory effects across human tissues. Science. 2020;369:1318–30.

16. Pan ZY, Wang Y, Wang MS, Wang YZ, Zhu XN, Gu SW, et al. An atlas of regulatory elements in chicken: A resource for chicken genetics and genomics. Science Advances. 2023;9:1204.

17. Clark EL, Archibald AL, Daetwyler HD, Groenen MAM, Harrison PW, Houston RD, et al. From FAANG to fork: application of highly annotated genomes to improve farmed animal production. Genome Biology. 2020;21:285.

18. Andersson L, Archibald AL, Bottema CD, Brauning R, Burgess SC, Burt DW, et al. Coordinated international action to accelerate genome-to-phenome with FAANG, the Functional Annotation of Animal Genomes project. Genome Biology. 2015;16:57.

19. Pan ZY, Yao YL, Yin HW, Cai ZX, Wang Y, Bai LJ, et al. Pig genome functional annotation enhances the biological interpretation of complex traits and human disease. Nature Communications. 2021;12:5848.

20. Liu SL, Gao YH, Canela-Xandri O, Wang S, Yu Y, Cai WT, et al. A multi-tissue atlas of regulatory variants in cattle. Nature Genetics. 2022;54:1438–47.

21. Teng J, Gao Y, Yin H, Bai Z, Liu S, Zeng H, et al. A compendium of genetic regulatory effects across pig tissues. Nature genetics. 2024;56:112–23.

22. Clark EL, Bush SJ, McCulloch MEB, Farquhar IL, Young R, Lefevre L, et al. A high resolution atlas of gene expression in the domestic sheep (Ovis aries). Plos Genetics. 2017;13:e1006997.

23. Davenport KM, Massa AT, Bhattarai S, McKay SD, Mousel MR, Herndon MK, et al. Characterizing genetic regulatory elements in ovine tissues. Frontiers in Genetics. 2021;12:628849.

24. Naval-Sanchez M, Nguyen Q, McWilliam S, Porto-Neto LR, Tellam R, Vuocolo T, et al. Sheep genome functional annotation reveals proximal regulatory elements contributed to the evolution of modern breeds. Nature Communications. 2018;9:859.

25. Bush SJ, McCulloch MEB, Muriuki C, Salavati M, Davis GM, Farquhar IL, et al. Comprehensive transcriptional profiling of the gastrointestinal tract of ruminants from birth to adulthood reveals strong developmental stage specific gene expression. G3-Genes Genomes Genetics. 2019;9:359–73.

26. Pokharel K, Peippo J, Weldenegodguad M, Honkatukia M, Li MH, Kantanen J. Gene expression profiling of corpus iuteum reveals important insights about early pregnancy in domestic sheep. Genes. 2020;11:415.

27. Arora R, Siddaraju NK, Manjunatha SS, Sudarshan S, Fairoze MN, Kumar A, et al. Muscle transcriptome provides the first insight into the dynamics of gene expression with progression of age in sheep. Scientific Reports. 2021;11:22360.

28. Zhao BR, Luo HP, He JM, Huang XX, Chen SQ, Fu XF, et al. Comprehensive transcriptome and methylome analysis delineates the biological basis of hair follicle development and wool-related traits in Merino sheep. Bmc Biology. 2021;19:197.

29. Pantalacci S, Semon M. Transcriptomics of developing embryos and organs: a raising tool for evo-devo. Journal of Experimental Zoology Part B-Molecular and Developmental Evolution. 2015;324:363–71.

30. Cao JY, Spielmann M, Qiu XJ, Huang XF, Ibrahim DM, Hill AJ, et al. The single-cell transcriptional landscape of mammalian organogenesis. Nature. 2019;566:496–502.

31. Cardoso-Moreira M, Halbert J, Valloton D, Velten B, Chen C, Shao Y, et al. Gene expression across mammalian organ development. Nature. 2019;571:505–09.

32. Valasi I, Barbagianni MS, Ioannidi KS, Vasileiou NGC, Fthenakis GC, Pourlis A. Developmental anatomy of sheep embryos, as assessed by means of ultrasonographic evaluation. Small Ruminant Research. 2017;152:56–73.

33. Gharibeh L, Yamak A, Whitcomb J, Lu AZ, Joyal M, Komati H, Liang WB, Fiset C, Nemer M. GATA6 is a regulator of sinus node development and heart rhythm. Proceedings of the National Academy of Sciences of the United States of America. 2021;118:e2007322118.

34. Webb EA, AlMutair A, Kelberman D, Bacchelli C, Chanudet E, Lescai F, et al. ARNT2 mutation causes hypopituitarism, post-natal microcephaly, visual and renal anomalies. Brain. 2013;136:3096–105.

35. Luo W, Li GH, Yi ZH, Nie QH, Zhang XQ. E2F1-miR-20a-5p/20b-5p auto-regulatory feedback loop involved in myoblast proliferation and differentiation. Scientific Reports. 2016;6:27904.

36. Moreno CS. SOX4: The unappreciated oncogene. Seminars in Cancer Biology. 2020;67:57–64.

37. Wang J, Huang YZ, Xu JW, Yue BL, Wen YF, Wang X, Lei CZ, Chen H. Pleomorphic adenoma gene 1 (PLAG1) promotes proliferation and inhibits apoptosis of bovine primary myoblasts through the PI3K-Akt signaling pathway. Journal of Animal Science. 2022;100:skac098.

38. Wyzykowski JC, Winata TI, Mitin N, Taparowsky EJ, Konieczny SF. Identification of novel MyoD gene targets in proliferating myogenic stem cells. Molecular and Cellular Biology. 2002;22:6199–208.

39. Gao SJ, Huang S, Zhang YH, Fang GM, Liu Y, Zhang CC, Li YL, Du J. The transcriptional regulator KLF15 is necessary for myoblast differentiation and muscle regeneration by activating FKBP5. Journal of Biological Chemistry. 2023;299:105226.

40. Pingault V, Zerad L, Bertani-Torres W, Bondurand N. SOX10: 20 years of phenotypic plurality and current understanding of its developmental function. Journal of Medical Genetics. 2022;59:105–14.

41. Hayashi S, Manabe I, Suzuki Y, Relaix F, Oishi Y. Klf5 regulates muscle differentiation by directly targeting muscle-specific genes in cooperation with MyoD in mice. Elife. 2016;5:e17462.

42. Min IM, Pietramaggiori G, Kim FS, Passegue E, Stevenson KE, Wagers AJ. The transcription factor EGR1 controls both the proliferation and localization of hematopoietic stem cells. Cell Stem Cell. 2008;2:380–91.

43. Liang T, Chen JR, Xu GY, Zhang ZD, Xue J, Zeng HP, et al. STAT3 and SPI1, may lead to the immune system dysregulation and heterotopic ossification in ankylosing spondylitis. Bmc Immunology. 2022;23:3.

44. Tidball JG. Regulation of muscle growth and regeneration by the immune system. Nature Reviews Immunology. 2017;17:165–78.

45. Fu X, Zhuang C-l, Hu P. Regulation of muscle stem cell fate. Cell Regeneration. 2022;11:40.

46. Zeitvogel J, Jokmin N, Rieker S, Klug I, Brandenberger C, Werfel T. GATA3 regulates FLG and FLG2 expression in human primary keratinocytes. Scientific Reports. 2017;7:11847.

47. Blum R, Vethantham V, Bowman C, Rudnicki M, Dynlacht BD. Genome-wide identification of enhancers in skeletal muscle: the role of MyoD1. Genes & Development. 2012;26:2763–79.

48. Packialakshmi B, Hira S, Lund K, Zhang AH, Halterman J, Feng YY, Scott DW, Lees JR, Zhou XM. NFAT5 contributes to the pathogenesis of experimental autoimmune encephalomyelitis (EAE) and decrease of T regulatory cells in female mice. Cellular Immunology. 2022;375:104515.

49. Tapias A, Ciudad CJ, Roninson IB, Noe V. Regulation of Sp1 by cell cycle related proteins. Cell Cycle. 2008;7:2856–67.

50. Vasu VT, Cross CE, Gohil K. Nrldl, an important circadian pathway regulatory gene, is suppressed by cigarette smoke in murine lungs. Integrative Cancer Therapies. 2009;8:321–28.

51. Hunter AL, Pelekanou CE, Barron NJ, Northeast RC, Grudzien M, Adamson AD, et al. Adipocyte NR1D1 dictates adipose tissue expansion during obesity. Elife. 2021;10.

52. Ka NL, Park MK, Kim SS, Jeon Y, Hwang S, Kim SM, Lim GY, Lee H, Lee MO. NR1D1 stimulates antitumor immune responses in breast cancer by activating cGAS-STING signaling. Cancer Research. 2023;83:3045–58.

53. Zhang Y, Xu S, Fan M, Yao H, Jiang C, He Q, Shi H, Lin R. Circadian rhythm disruption modulates enteric neural precursor cells differentiation leading to gastrointestinal motility dysfunction via the NR1D1/NF-kappaB axis. Journal of translational medicine. 2024;22:975.

54. Langfelder P, Horvath S. WGCNA: an R package for weighted correlation network analysis. Bmc Bioinformatics. 2008;9:559.

55. Lemoine GG, Scott-Boyer MP, Ambroise B, Perin O, Droit A. GWENA: gene co-expression networks analysis and extended modules characterization in a single Bioconductor package. Bmc Bioinformatics. 2021;22:267.

56. Hyvarinen A, Oja E. Independent component analysis: algorithms and applications. Neural Networks. 2000;13:411–30.

57. Stegle O, Parts L, Piipari M, Winn J, Durbin R. Using probabilistic estimation of expression residuals (PEER) to obtain increased power and interpretability of gene expression analyses. Nature Protocols. 2012;7:500–07.

58. Song WM, Zhang B. Multiscale Embedded Gene Co-expression Network Analysis. Plos Computational Biology. 2015;11:e1004574.

59. Russo PST, Ferreira GR, Cardozo LE, Burger MC, Arias-Carrasco R, Maruyama SR, et al. CEMiTool: a Bioconductor package for performing comprehensive modular co-expression analyses. Bmc Bioinformatics. 2018;19:56.

60. Stein-O’Brien GL, Arora R, Culhane AC, Favorov AV, Garmire LX, Greene CS, et al. Enter the matrix: factorization uncovers knowledge from omics. Trends in Genetics. 2018;34:790–805.

61. Way GP, Zietz M, Rubinetti V, Himmelstein DS, Greene CS. Compressing gene expression data using multiple latent space dimensionalities learns complementary biological representations. Genome Biology. 2020;21.

62. Lazure F, Blackburn DM, Corchado AH, Sahinyan K, Karam N, Sharanek A, et al. Myf6/MRF4 is a myogenic niche regulator required for the maintenance of the muscle stem cell pool. Embo Reports. 2020;21:e49499.

63. Ganassi M, Badodi S, Wanders K, Zammit PS, Hughes SM. Myogenin is an essential regulator of adult myofibre growth and muscle stem cell homeostasis. Elife. 2020;9:e60445.

64. Wang YK, Schnegelsberg PNJ, Dausman J, Jaenisch R. Functional redundancy of the muscle-specific transcription factors Myf5 and myogenin. Nature. 1996;379:823–25.

65. Milan G, Romanello V, Pescatore F, Armani A, Paik JH, Frasson L, et al. Regulation of autophagy and the ubiquitin-proteasome system by the FoxO transcriptional network during muscle atrophy. Nature Communications. 2015;6:6670.

66. Xu M, Chen XL, Chen DW, Yu B, Huang ZQ. FoxO1: a novel insight into its molecular mechanisms in the regulation of skeletal muscle differentiation and fiber type specification. Oncotarget. 2017;8:10662–74.

67. Ram PA, Park SH, Choi HK, Waxman DJ. Growth hormone activation of Stat 1, Stat 3, and Stat 5 in rat liver - differential kinetics of hormone desensitization and growth hormone stimulation of both tyrosine phosphorylation and serine/threonine phosphorylation. Journal of Biological Chemistry. 1996;271:5929–40.

68. Mallipattu SK, Estrada CC, He JC. The critical role of Kruppel-like factors in kidney disease. American Journal of Physiology-Renal Physiology. 2017;312:F259–F65.

69. Sobiak B, Graczyk A, Lesniak W. DNA methylation analysis of selected genes comprised in the epidermal differentiation complex (EDC) during epidermal differentiation. Febs Journal. 2014;281:300–00.

70. Kypriotou M, Huber M, Hohl D. The human epidermal differentiation complex: cornified envelope precursors, S100 proteins and the ‘fused genes’ family. Experimental Dermatology. 2012;21:643–49.

71. Abhishek S, Krishnan SP. Epidermal differentiation complex: a review on its epigenetic regulation and potential drug targets. Cell Journal. 2016;18:1–6.

72. Lenffer J, Nicholas FW, Castle K, Rao A, Gregory S, Poidinger M, Mailman MD, Ranganathan S. OMIA (Online Mendelian Inheritance in Animals): an enhanced platform and integration into the Entrez search interface at NCBI. Nucleic Acids Research. 2006;34:D599–D601.

73. Cooper TA. Regulating mRNA complexity in the mammalian brain. Nature Genetics. 2011;43:618–19.

74. Wright CJ, Smith CWJ, Jiggins CD. Alternative splicing as a source of phenotypic diversity. Nature Reviews Genetics. 2022;23:697–710.

75. Naro C, Cesari E, Sette C. Splicing regulation in brain and testis: common themes for highly specialized organs. Cell Cycle. 2021;20:480–89.

76. Jin L, Tang QZ, Hu SL, Chen ZX, Zhou XM, Zeng B, et al. A pig BodyMap transcriptome reveals diverse tissue physiologies and evolutionary dynamics of transcription. Nature Communications. 2021;12.

77. Cabili MN, Trapnell C, Goff L, Koziol M, Tazon-Vega B, Regev A, Rinn JL. Integrative annotation of human large intergenic noncoding RNAs reveals global properties and specific subclasses. Genes & Development. 2011;25:1915–27.

78. Dailu Guan, Zhonghao Bai, Xiaoning Zhu, Conghao Zhong, Yali Hou, The ChickenGTEx Consortium, et al. The ChickenGTEx pilot analysis: a reference of regulatory variants across 28 chicken tissues. bioRxiv. 2023.

79. Yan Z, Yang J, Wei WT, Zhou ML, Mo DX, Wan X, et al. A time-resolved multi-omics atlas of transcriptional regulation in response to high-altitude hypoxia across whole-body tissues. Nature Communications. 2024;15:3970.

80. Li C, Chen BC, Langda S, Pu P, Zhu XJ, Zhou SW, et al. Multi-omic analyses shed light on the genetic control of high-altitude adaptation in sheep. Genomics Proteomics & Bioinformatics. 2024;22:qzae030.

81. Vidal VPI, Chaboissier MC, Lützkendorf S, Cotsarelis S, Mill P, Hui CC, Ortonne N, Ortonne JP, Schedl A. Sox9 is essential for outer root sheath differentiation and the formation of the hair stem cell compartment. Current Biology. 2005;15:1340–51.

82. Dall’Olio S, Fontanesi L, Tognazzi L, Russo V. Genetic structure of candidate genes for litter size in Italian Large White pigs. Veterinary Research Communications. 2010;34:S203–S06.

83. An XP, Han D, Hou JX, Li G, Wang JG, Yang MM, et al. GnRHR gene polymorphisms and their effects on reproductive performance in Chinese goats. Small Ruminant Research. 2009;85:130–34.

84. Horodyska J, Hamill RM, Varley PF, Reyer H, Wimmers K. Genome-wide association analysis and functional annotation of positional candidate genes for feed conversion efficiency and growth rate in pigs. Plos One. 2017;12.

85. Kominakis A, Hager-Theodorides AL, Zoidis E, Saridaki A, Antonakos G, Tsiamis G. Combined GWAS and ‘guilt by association’-based prioritization analysis identifies functional candidate genes for body size in sheep. Genetics Selection Evolution. 2017;49.

86. Feng W, Bais A, He HT, Rios C, Jiang S, Xu J, Chang C, Kostka D, Li G. Single-cell transcriptomic analysis identifies murine heart molecular features at embryonic and neonatal stages. Nature Communications. 2022;13:7960.

87. Mantri M, Scuderi GJ, Abedini-Nassab R, Wang MFZ, McKellar D, Shi H, Grodner B, Butcher JT, De Vlaminck I. Spatiotemporal single-cell RNA sequencing of developing chicken hearts identifies interplay between cellular differentiation and morphogenesis. Nature Communications. 2021;12:1771.

88. Wang F, Ding PW, Liang X, Ding XN, Brandt CB, Sjostedt E, et al. Endothelial cell heterogeneity and microglia regulons revealed by a pig cell landscape at single-cell level. Nature Communications. 2022;13:3620.

89. Li HY, Zhou JX, Li ZX, Chen SY, Liao XY, Zhang B, et al. A comprehensive benchmarking with practical guidelines for cellular deconvolution of spatial transcriptomics. Nature Communications. 2023;14:1548.

90. Holler K, Neuschulz A, Drewe-Boss P, Mintcheva J, Spanjaard B, Arsie R, Ohler U, Landthaler M, Junker JP. Spatio-temporal mRNA tracking in the early zebrafish embryo. Nature Communications. 2021;12:3358.

91. Lagarde J, Uszczynska-Ratajczak B, Carbonell S, Perez-Lluch S, Abad A, Davis C, et al. High-throughput annotation of full-length long noncoding RNAs with capture long-read sequencing. Nature Genetics. 2017;49:1731–40.

92. He P, Williams BA, Trout D, Marinov GK, Amrhein H, Berghella L, et al. The changing mouse embryo transcriptome at whole tissue and single-cell resolution. Nature. 2020;583:760–67.

93. Sarropoulos I, Marin R, Cardoso-Moreira M, Kaessmann H. Developmental dynamics of lncRNAs across mammalian organs and species. Nature. 2019;571:510.

94. Bolger AM, Lohse M, Usadel B. Trimmomatic: a flexible trimmer for Illumina sequence data. Bioinformatics. 2014;30:2114–20.

95. Dobin A, Davis CA, Schlesinger F, Drenkow J, Zaleski C, Jha S, Batut P, Chaisson M, Gingeras TR. STAR: ultrafast universal RNA-seq aligner. Bioinformatics. 2013;29:15–21.

96. Liao Y, Smyth GK, Shi W. featureCounts: an efficient general purpose program for assigning sequence reads to genomic features. Bioinformatics. 2014;30:923–30.

97. Kovaka S, Zimin AV, Pertea GM, Razaghi R, Salzberg SL, Pertea M. Transcriptome assembly from long-read RNA-seq alignments with StringTie2. Genome Biology. 2019;20:278.

98. Patro R, Duggal G, Love MI, Irizarry RA, Kingsford C. Salmon provides fast and bias-aware quantification of transcript expression. Nature Methods. 2017;14:417–19.

99. Bray NL, Pimentel H, Melsted P, Pachter L. Near-optimal probabilistic RNA-seq quantification. Nature Biotechnology. 2016;34:525–27.

100. Li YI, Knowles DA, Humphrey J, Barbeira AN, Dickinson SP, Im HK, Pritchard JK. Annotation-free quantification of RNA splicing using LeafCutter. Nature Genetics. 2018;50:151–58.

101. Camargo AP, Vasconcelos AA, Fiamenghi MB, Pereira GAG, Carazzolle MF. tspex: a tissue-specificity calculator for gene expression data. Research Square (2020). 2020; doi:10.21203/rs.3.rs-51998/v1.

102. Robinson MD, McCarthy DJ, Smyth GK. edgeR: a Bioconductor package for differential expression analysis of digital gene expression data. Bioinformatics. 2010;26:139–40.

103. Wu T, Hu E, Xu S, Chen M, Guo P, Dai Z, et al. clusterProfiler 4.0: A universal enrichment tool for interpreting omics data. Innovation (New York, NY). 2021;2:100141.

104. Hanzelmann S, Castelo R, Guinney J. GSVA: gene set variation analysis for microarray and RNA-Seq data. Bmc Bioinformatics. 2013;14:7.

105. Bailey TL, Boden M, Buske FA, Frith M, Grant CE, Clementi L, Ren JY, Li WW, Noble WS. MEME SUITE: tools for motif discovery and searching. Nucleic Acids Research. 2009;37:W202–W08.

106. Fornes O, Castro-Mondragon JA, Khan A, van der Lee R, Zhang X, Richmond PA, et al. JASPAR 2020: update of the open-access database of transcription factor binding profiles. Nucleic Acids Research. 2020;48:D87–D92.

107. Kulakovskiy IV, Vorontsov IE, Yevshin IS, Sharipov RN, Fedorova AD, Rumynskiy EI, et al. HOCOMOCO: towards a complete collection of transcription factor binding models for human and mouse via large-scale ChIP-Seq analysis. Nucleic Acids Research. 2018;46:D252–D59.

108. Futschik ME, Carlisle B. Noise-robust soft clustering of gene expression time-course data. Journal of Bioinformatics and Computational Biology. 2005;3:965–88.

109. Kumar L, E Futschik M. Mfuzz: a software package for soft clustering of microarray data. Bioinformation. 2007;2:5–7.

110. Leonard M, Graham S, Bonacum D. The human factor: the critical importance of effective teamwork and communication in providing safe care. Quality & Safety in Health Care. 2004;13:I85–I90.

111. Chung NC, Miasojedow B, Startek M, Gambin A. Jaccard/Tanimoto similarity test and estimation methods for biological presence-absence data. Bmc Bioinformatics. 2019;20:644.

112. Langfelder P, Luo R, Oldham MC, Horvath S. Is My Network Module Preserved and Reproducible? Plos Computational Biology. 2011;7:e1001057.

113. Xue ZG, Huang K, Cai CC, Cai LB, Jiang CY, Feng Y, et al. Genetic programs in human and mouse early embryos revealed by single-cell RNA sequencing. Nature. 2013;500:593–97.

114. Browning BL, Zhou Y, Browning SR. A one-penny imputed genome from next-generation reference panels. American Journal of Human Genetics. 2018;103:338–48.

115. Yang JA, Lee SH, Goddard ME, Visscher PM. GCTA: a tool for genome-wide complex trait analysis. American Journal of Human Genetics. 2011;88:76–82.

116. Nelson CP, Goel A, Butterworth AS, Kanoni S, Webb TR, Marouli E, et al. Association analyses based on false discovery rate implicate new loci for coronary artery disease. Nature Genetics. 2017;49:1385–91.

117. Purcell S, Neale B, Todd-Brown K, Thomas L, Ferreira MAR, Bender D, et al. PLINK: A tool set for whole-genome association and population-based linkage analyses. American Journal of Human Genetics. 2007;81:559–75.

118. Wang G, Sarkar A, Carbonetto P, Stephens M. A simple new approach to variable selection in regression, with application to genetic fine mapping. Journal of the Royal Statistical Society Series B-Statistical Methodology. 2020;82:1273–300.

119. Rohde PD, Fourie Sorensen I, Sorensen P. qgg: an R package for large-scale quantitative genetic analyses. Bioinformatics. 2020;36:2614–15.

120. Xiang RD, McNally J, Bond J, Tucker D, Cameron M, Donaldson AJ, et al. Across-experiment transcriptomics of sheep rumen identifies expression of lipid/oxo-acid metabolism and muscle cell junction genes associated with variation in methane-related phenotypes. Frontiers in Genetics. 2018;9:330.

